# ARF1 compartments direct cargo flow via maturation into recycling endosomes

**DOI:** 10.1101/2023.10.27.564143

**Authors:** Alexander Stockhammer, Petia Adarska, Vini Natalia, Anja Heuhsen, Antonia Klemt, Gresy Bregu, Shelly Harel, Carmen Rodilla-Ramirez, Carissa Spalt, Ece Özsoy, Paula Leupold, Alica Grindel, Eleanor Fox, Joy Orezimena Mejedo, Amin Zehtabian, Helge Ewers, Dmytro Puchkov, Volker Haucke, Francesca Bottanelli

## Abstract

Cellular membrane homeostasis is maintained via a tightly regulated membrane and cargo flow between organelles of the endocytic and secretory pathways. Adaptor protein complexes (APs), which are recruited to membranes by the small GTPase ARF1, facilitate cargo selection and incorporation into trafficking intermediates. According to the classical model, small vesicles would facilitate bi-directional long-range transport between the Golgi, endosomes and plasma membrane. Here we revisit the intracellular organization of the vesicular transport machinery using a combination of CRISPR-Cas9 gene editing, live-cell high temporal (fast-confocal) or spatial (stimulated emission depletion (STED)) microscopy as well as correlative light and electron microscopy. We characterize novel tubulo-vesicular ARF1 compartments that harbor clathrin and different APs. Our findings reveal two functionally different classes of ARF1 compartments, each decorated by a different combination of APs. Perinuclear ARF1 compartments facilitate Golgi export of secretory cargo, while peripheral ARF1 compartments are involved in endocytic recycling downstream of early endosomes. Contrary to the classical model of long-range vesicle shuttling, we observe that ARF1 compartments shed ARF1 and mature into recycling endosomes. This maturation process is impaired in the absence of AP-1 and results in trafficking defects. Collectively, these data highlight a crucial role for ARF1 compartments in post-Golgi sorting.

## Introduction

Eucaryotic cells compartmentalize biochemical reactions within membrane bound organelles. Material exchange occurs either via transport carriers that bud from a donor- and fuse with an acceptor-compartment or via direct contact between membranes. The Golgi apparatus and endosomal organelles are crucial for maintaining cellular membrane homeostasis as they coordinate the busiest trafficking highways at the intersection between exocytic and endocytic traffic. According to current models, communication between the Golgi and various endosomes is mediated by clathrin-coated vesicles^1,2^. Vesicle formation is orchestrated by a complex protein machinery, which ensures the specificity of cargo selection and guides trafficking intermediates to their correct destination within the cell. For this, key cargo adaptors such as the adaptor protein complex 1 (AP-1) are recruited to membranes by the ADP-ribosylation factor 1 (ARF1)^2–4^, where they can recruit cargo and coat proteins such as clathrin to drive membrane curvature and vesicle formation^1,5^.

When imaged as fusion proteins or with antibodies, clathrin and adaptors define punctate structures throughout the cytoplasm of the cell, supporting a vesicular model for cargo exchange between the Golgi, endosomes and the plasma membrane^5–8^. In contrast to shuttling vesicles and full-collapse fusion events, intracellular organelle communication through a kiss-and-run mechanism was proposed^9^. Various secretory and endocytic recycling cargoes have been found to transit through tubulo-vesicular compartments^10,11^, some of which were shown to be decorated with clathrin and AP-1^12–14^. In particular, secretory cargo flow from the Golgi to the plasma membrane may occur via direct tubular carriers or clathrin-decorated tubules that deliver their content to endosomes via yet unknown mechanisms^13,15^. Tubulo-vesicular endosomes harboring the adaptors AP-1 and AP-3 as well as clathrin have been identified using immuno-EM but have not been characterized in detail^16,17^. Cargo sorting from sorting endosomal compartments was also shown to occur via formation of tubular domain responsible for cargo sequestration^18^. At early endosomes, cargo enrichment and tubulation of the membrane depend on sorting nexins and sorting complexes such as retromer, retriever and the CCC complex^19^. This process is important to sequester cargoes destined for recycling to the plasma membrane and the Golgi away from early endosomes, which subsequently mature into late endosomes and finally fuse with lysosomes to delivery their content^20–22^.

Investigating the mechanisms and dynamics of proteins sorting out of the Golgi and within the endo-lysosomal system remains challenging, urging a combination of approaches aimed at investigating dynamic events and the underlying ultrastructure in near-physiological conditions. First, live-cell imaging is indispensable not only because of the transient and highly dynamic nature of many trafficking events, but also because post-Golgi structures, such as tubulo-vesicular compartments, are lost upon fixation^23^. Second, overexpression of fusion proteins, which is often used for making proteins accessible for live-cell imaging, holds the potential for artifacts^24^, as too high protein levels can affect protein localization and function, making endogenous tagging crucial. Third, visualization of sorting events has proven to be a difficult task in the perinuclear area where organelles are tightly packed, demanding super-resolution imaging techniques suitable for imaging live specimens^13,25,26^. Here, we utilize CRISPR-Cas9 technology to endogenously tag various sorting machinery components involved in post-Golgi trafficking with the self-labeling enzymes HaloTag and SNAP-tag^27^. Employing various imaging techniques, such as 3D correlative light electron microscopy, fast live-cell confocal and super-resolution STED microscopy, we were able to characterize novel multi-functional tubulo-vesicular ARF1 sorting compartment harboring different adaptor proteins and clathrin. Trafficking assays and CRISPR-Cas9 mediated knock-outs point at a role for ARF1 compartments in endocytic recycling (downstream of early endosomes marked by Rab5) and post-Golgi secretory traffic. Interestingly, for both secretory and endocytic trafficking, cargo is observed first in ARF1 compartments and subsequently in recycling endosomes (REs). This prompted us to investigate how cargo is transferred from ARF1 compartments to REs. We show that ARF1 compartments undergo maturation into REs via a mechanism that depends on AP-1, as loss of AP-1 inhibits maturation and causes trafficking defects. Advancing previous thinking in this field, our findings suggest a model where cargo sorting is mediated by a dynamic tubular network that connects the trans-Golgi network (TGN) with endo-lysosomes and the plasma membrane^20,28,29^.

## Results

### Clathrin is associated with ARF1 compartments and organizes their fission

Previous studies have identified TGN-derived tubulo-vesicular compartments defined by the small GTPase ARF1^13, 23^. Interestingly, these compartments were found to harbor clathrin nanodomains. This observation raised the possibility that they could be hubs for clathrin-coated vesicle budding and define a tubulo-vesicular network responsible for post-Golgi cargo sorting. First, we recapitulated our initial observations^23^ and we applied super-resolution live-cell STED microscopy on ARF1^EN^-Halo/SNAP-CLCa^EN^ (EN=endogenous) knock-in (KI) HeLa cells to highlight the close association of clathrin with ARF1 compartments (Fig. 1A). Additionally, we observed ARF1-positive clathrin compartments in various cell types (Extended Data Fig. 1A-B). To further characterize ARF1 compartments, we first wanted to test whether non-endocytic clathrin is exclusively associated with ARF1 compartments. For this, we created an ARF1^EN^-eGFP/Halo-CLCa^EN^/AP2µ^EN^-SNAP triple KI cell line (Fig. 1B). Quantification of clathrin association with either AP-2 (endocytic)^30^ or ARF1 revealed that most of the non-endocytic and non-Golgi clathrin decorates ARF1-positive membranes (Fig. 1C). Furthermore, we took advantage of fast confocal live-cell imaging to get a better understanding of the dynamics of clathrin on ARF1-compartments (Fig. 1D, Extended Data Video 1). Notably, we could not visualize any clathrin-coated vesicles budding from ARF1 compartments. Instead, we observed clathrin clusters translocating together with the closely associated membrane (Fig. 1E). Strikingly, when we looked at ARF1 compartments emerging from the TGN, we found clathrin associated with the detaching tubule (Fig. 1F). Without visualization of the underlying ARF1 membrane, these events could have easily been mistaken for a clathrin-coated vesicle moving within the cytoplasm or budding from the TGN. Intriguingly, clathrin localized at the fission site on ARF1 compartments, suggesting that clathrin and associated machinery may be responsible for the recruitment of fission factors. We analyzed more than 100 fission events and found that clathrin was present at >90% of the fission sites (Fig 1G-H, Extended Data Fig. 1C).

**Figure 1:**
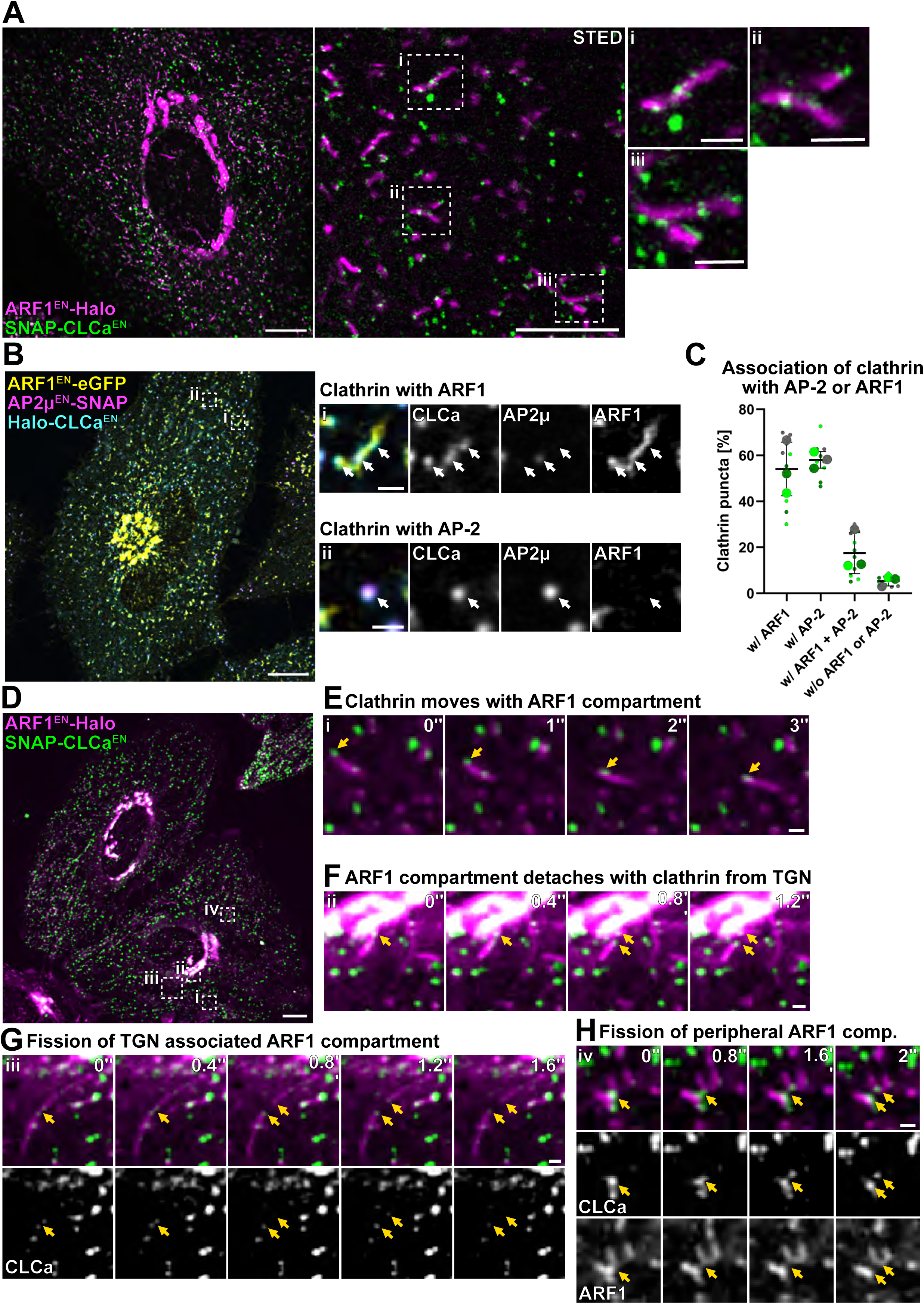
ARF1 compartments are the major site of non-endocytic clathrin assembly. (**A**) Live-cell confocal and STED imaging of ARF1^EN^-Halo/SNAP-CLCa^EN^ HeLa cells labeled with CA-JF_571_ and BG-JFX_650_, show association of clathrin to ARF1 compartments. (**B**) Live-cell confocal imaging of ARF1^EN^-eGFP/AP2µ^EN^-SNAP/Halo-CLCa^EN^ HeLa cells labeled with CA-JF_552_ and BG-JFX_650_, highlighting association of **(i)** non-endocytic clathrin with ARF1 compartments and **(ii)** endocytic clathrin with AP-2. (**C**) Quantification of clathrin-association with ARF1 and/or AP-2. In total 10 cells from 3 independent experiments were analyzed, replicates are shown in different colors and each small dot represents a single cell of the replicate. (**D**) Time-lapse confocal spinning disk imaging of ARF1^EN^-Halo/SNAP-CLCa^EN^ HeLa cells labeled with CA-JF_552_ and BG-JFX_650_, (**E**) highlights movement of clathrin together with ARF1 compartments and (**F**) detachment of ARF1 compartments from the TGN together with clathrin. (**G**) Clathrin is found at sites of fission of ARF1 compartments when they detach from the TGN and (**H**) in the cell periphery. Selected frames are shown, movie was taken with a frame rate of 5 frames/s. EN=endogenous, BG=benzylguanine (SNAP-tag substrate), CA=chloroalkane (HaloTag substrate). Scale bars: 10 µm (confocal overview), 5 µm (STED image in **A**) and 1 µm (crops).

To get a better understanding of the membrane organization of ARF1 compartments and the association of clathrin with these membranes, we employed three-dimensional correlative light electron microscopy (3D-CLEM). ARF1^EN^-Halo/SNAP-CLCa^EN^ KI cells were labeled with cell-permeable dyes and imaged post-fixation by scanning confocal microscopy (Fig. 2A), embedded, and a small region of interest was visualized using focused ion beam scanning electron microscopy (FIB-SEM) imaging (Fig. 2B). Alignment of confocal light microscopy and FIB-SEM imaging enabled the identification of ARF1 compartments at an isotropic resolution of 7 nm (Fig. 2Ci-iii, Extended Data Video 2). Intriguingly, the underlying tubulo-vesicular compartments displayed a pearled morphology that is reminiscent of the morphology of the ER-Golgi intermediate tubular compartments that mediate ER export^31^. In contrast to the clathrin-coated vesicular structures, which have a confined diameter between 70 and 90 nm, the diameter of the non-clathrin-coated ARF1 compartment varied between 20 nm and 180 nm (Fig. 2D). Clathrin-positive membranes seem to be preferably located at sites with high intrinsic curvature and are directly connected to the ARF1 compartment via a membranous neck (Fig. 2E). In summary, these data identify ARF1 compartments as the major site of clathrin recruitment. This finding motivated us to further characterize the identity and function of these compartments.

**Figure 2:**
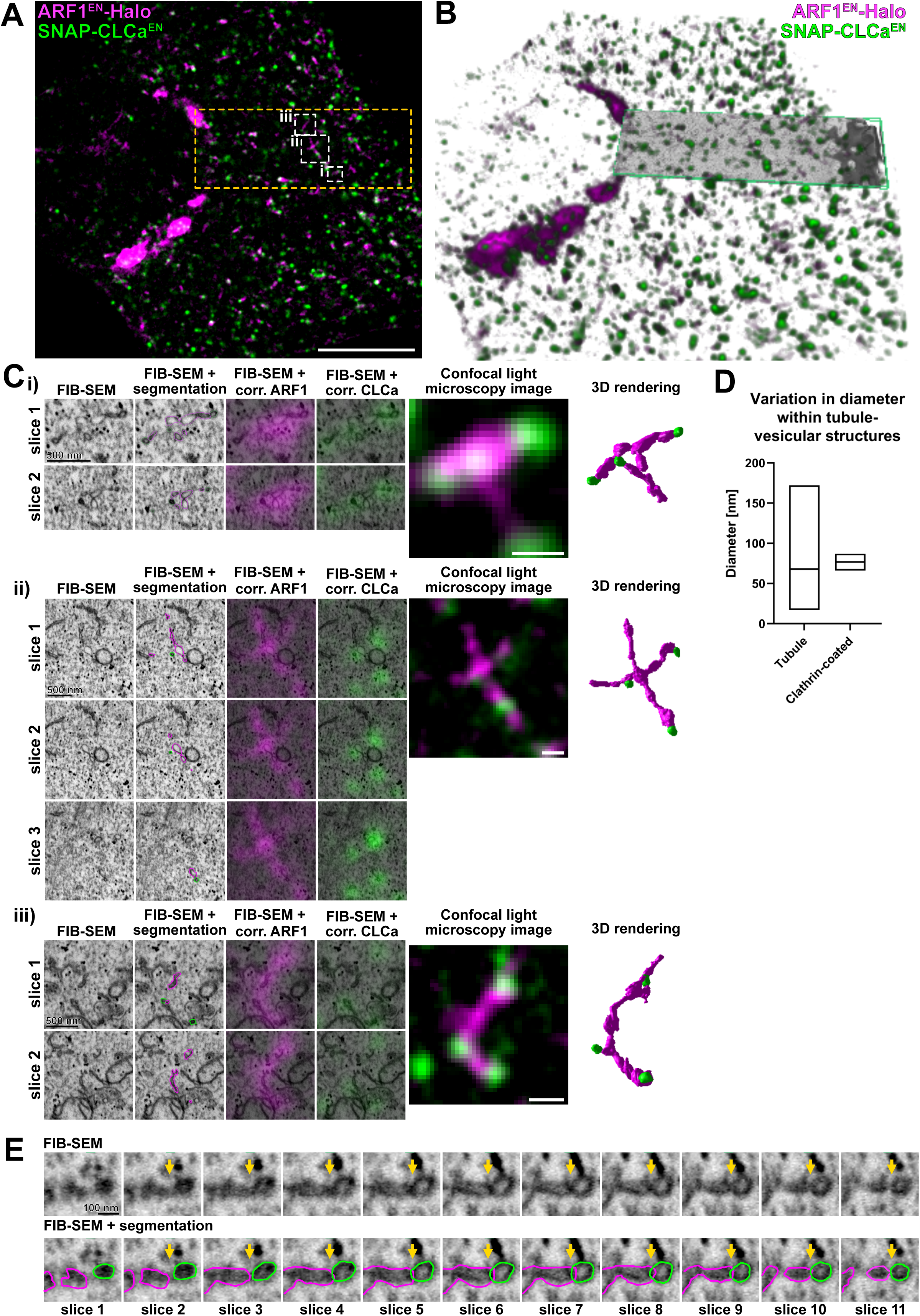
Tubulo-vesicular nature of ARF1 compartments revealed by 3D CLEM. (**A**) Slice of a confocal z-stack of ARF1^EN^-Halo/SNAP-CLCa^EN^ HeLa cell labeled with CA-JF_552_ and BG-JFX_650_ that was chosen for CLEM. The area that was imaged with FIB-SEM is highlighted with a yellow outline. Segmented ARF1 compartments **(i-iii)** are shown. (**B**) Overlay of the 3D projection of the confocal stack and FIB-SEM image (green box). FIB-SEM image was obtained with 7 nm isotropic resolution. (**C**) Segmentation of individual ARF1 compartments with clathrin. Shown are 2-3 exemplary slices of the FIB-SEM image, the FIB-SEM image with outlines from the segmented area, overlays of fluorescence of ARF1 and clathrin with the FIB-SEM image, a representative slice of the confocal image of the ARF1 compartments and the 3D rendering from the ARF1 compartments. (**D**) Diagram showing variation in tubule diameter of the ARF1 compartment and the clathrin coated areas. Data for the diagram was obtained from the three ARF1 compartments shown in **C** that were measured at different parts of the tubule. (**E**) Single slices (7 nm increments) of the FIB-SEM data set, including segmentation, highlight the connection of clathrin-coated and non-coated part of the ARF1 compartment (arrow highlights neck of non-clathrin coated and clathrin coated ARF1 compartment). EN=endogenous, BG=benzylguanine (SNAP-tag substrate), CA=chloroalkane (HaloTag substrate). Scale bars: 10 µm (overview in **A**), and 500 nm (crops in **C**).

### AP-1 and AP-3 localize to segregated nanodomains on ARF1 compartments, clathrin is recruited to AP-1 (but not AP-3) positive nanodomains

Post-Golgi protein sorting relies on the recruitment of different cargo adaptors, such as AP-1 or AP-3 to membranes. Both adaptors were previously reported to localize to the TGN^5,6,14^ and endosomal membranes^12,17,32^, where they coordinate bi-directional transport between TGN and endosomes (AP-1)^1,33,34^ or transport to late endosomes and melanosomes (AP-3)^17,32^. As both adaptors are recruited by ARF1^3,4^, we wondered whether they are present on ARF1 compartments. To test the localization of AP-1 and AP-3, we introduced a SNAP-tag to the C-terminus of their µA-subunit in ARF1^EN^-Halo KI cells and created double KI cell lines (Fig. 3A-B). Interestingly, as observed for clathrin (Fig. 1A), live-cell STED highlighted nanodomains of both AP-1 and AP-3 on ARF1 compartments (Fig. 3A-B). We observed the same localization pattern when we endogenously tagged the large AP1γ1- or AP3δ1-subunit (Extended Data Fig. 2A-B) suggesting that the placement of the tag within the complex does not impact AP localization. Furthermore, simultaneous tagging and visualization of medium and large subunits of AP complexes showed both subunits to colocalize within the same nanodomains, indicating that AP complexes still form when one or more subunits are tagged (Extended Data Fig. 2C-D). In addition, AP-1-dependent clathrin recruitment was unaffected by Halo-tagging of the µ-subunit (Extended Data Fig. 2E-F), indicating that interaction of AP-1 with accessory proteins is unaffected by addition of the imaging tags.

**Figure 3:**
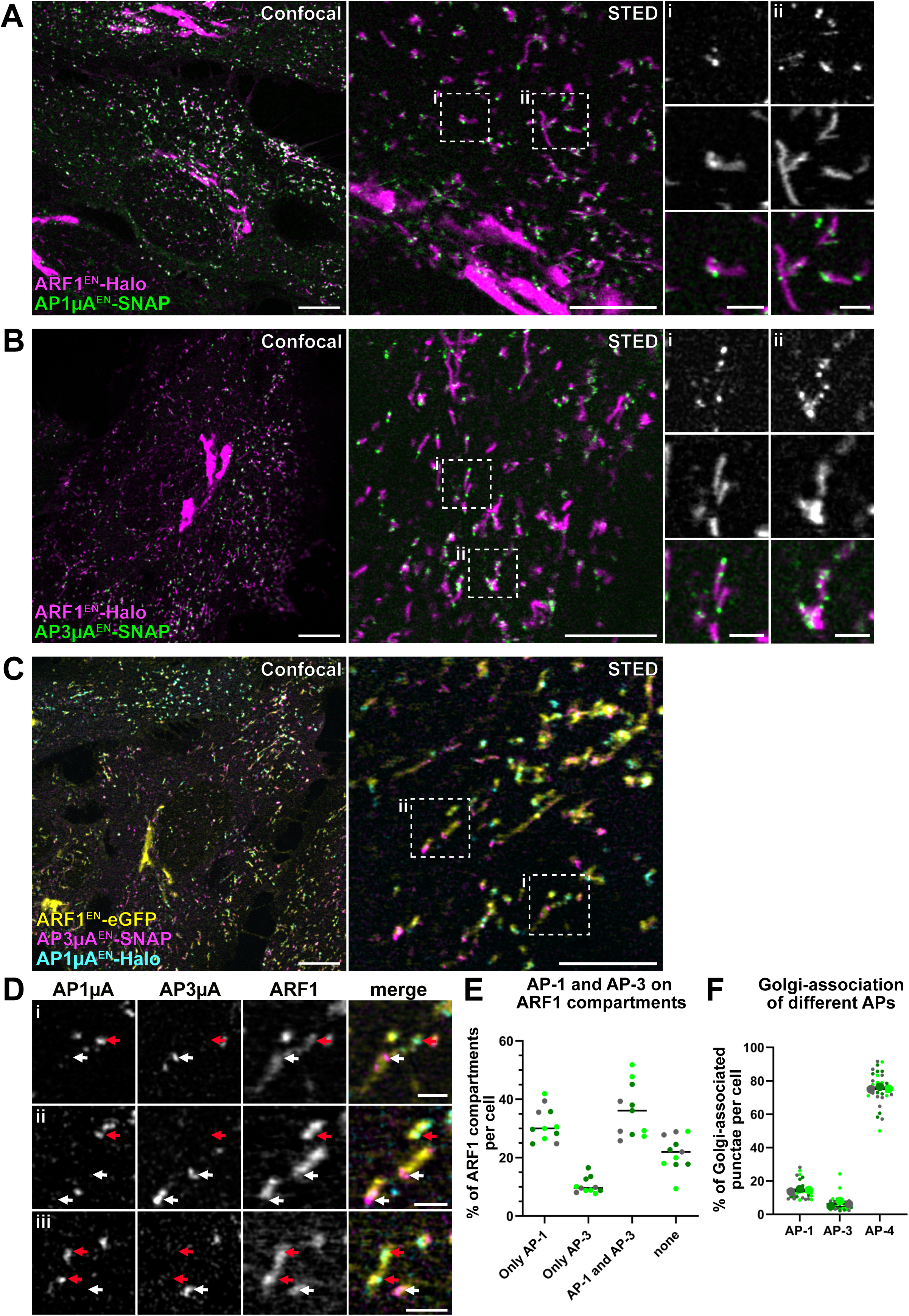
Adaptor protein complexes AP-1 and AP-3 define segregated nanodomains on ARF1 compartments. (**A-B**) Live-cell confocal and STED imaging of ARF1^EN^-Halo/AP1µA^EN^-SNAP HeLa cells (**A**) and ARF1^EN^-Halo/AP3µA^EN^-SNAP HeLa cells (**B**) labeled with CA-JF_571_ and BG-JFX_650_ show association of AP-1 and AP-3 to ARF1 compartments. **(C-D)** Live-cell confocal and STED imaging (2 color STED imaging with ARF1^EN^-eGFP imaged in confocal mode) of ARF1^EN^-eGFP/AP3µA^EN^-SNAP/AP1µA^EN^-Halo HeLa cells labeled with CA-JF_571_ and BG-JFX_650_ show that AP-1 and AP-3 localize to segregated nanodomains on ARF1 compartments. **(i)** Crops highlight that AP-1 (red arrows) and AP-3 (white arrows) localize to segregated nanodomains on the same compartment. **(ii-iii)** In addition, ARF1 compartments harboring either AP-1 or AP-3 are found. (**E**) Quantification of the percentage of ARF1 compartments with specific adaptor identity per cell. In total 11 cells from 3 independent experiments were analyzed, replicates are shown in different colors each dot representing a single cell. (**F**) Quantification of Golgi-associated puncta positive for AP-1, AP-3 and AP-4. In total 30 cells from 3 independent experiments were analyzed for each condition, replicates are shown in different colors and each small dot represents a single cell of the replicate. EN=endogenous, BG=benzylguanine (SNAP-tag substrate), CA=chloroalkane (HaloTag substrate). Scale bars: 10 µm (confocal overview), 5 µm (STED images) and 1 µm (STED crops).

As we observed both adaptors on ARF1 compartments, we asked whether these compartments represent multi-functional sorting endosomes responsible for both AP-1- and AP-3-dependent sorting or whether different classes of ARF1 compartments are defined by different adaptors. To understand the nature and function of the compartments, we created an ARF1^EN^-eGFP/AP1µA^EN^-Halo/AP3µA^EN^-SNAP triple KI cell line (Fig. 3C). Live-cell confocal and STED imaging revealed that both adaptors localize to segregated nanodomains on ARF1 compartments. The most abundant class of ARF1 compartments is decorated with AP-1 and AP-3 (∼38%) (Fig. 3C, Di, E). However, we could additionally observe ARF1 compartments that only supported AP-1 (∼30%) or AP-3 recruitment (∼10%) (Fig. 3Dii-iii, E). In particular, ARF1 compartments that were seen in the perinuclear area and emerging from the Golgi were only positive for AP-1. In contrast, tubules observed in the cell periphery were found to be positive for both AP-1 and AP-3 (Fig. 3A-C). These data suggest that functionally distinct populations of ARF1 compartments may co-exist. About 20% of all ARF1 compartments did not harbor any AP-1 or AP-3, in agreement with a role for ARF1 tubules in retrograde Golgi-to-ER transport^13,23^. Both AP-1 and AP-3 were often shown to localize predominantly to the TGN/Golgi area in fixed cells^5,6^. However, in living gene-edited cells we found that only around 16% of total AP-1 and 6% of total AP-3 punctae were confined to the perinuclear area (Fig. 3F). Fixation is known to disrupt tubulo-vesicular cytoplasmic membranes^23^ possibly leading to an overestimation of the fraction of Golgi-associated adaptors. Additionally, while a population of AP-1 punctae associated with the Golgi was very prominent, AP-3 was mainly excluded from the Golgi area (Fig. 3F). AP-4, the other adaptor protein complex recruited by ARF1^23,35^, was predominantly associated with the Golgi (Fig. 3F, Extended Data Fig. 2G). These data suggest that ARF1 compartments serve as the main hub for AP-1 and AP-3-dependend sorting and might fulfill distinct intracellular functions based on the adaptors present.

Next, we wanted to investigate whether both AP-1 and AP-3 have the ability to recruit clathrin in living cells. Clathrin binding to AP-1 is well established^36^. Although AP-3 was shown to bind clathrin *in vitro*^6^, the interaction of AP-3 with clathrin in mammalian cells is debated^17,37–39^. To resolve this issue, we created triple KI cell lines that allowed simultaneous visualization of ARF1, clathrin and AP-1 (ARF1^EN^-eGFP/AP1µA^EN^-SNAP/Halo-CLCa^EN^) or AP-3 (ARF1^EN^-eGFP/AP3µA^EN^-SNAP/Halo-CLCa^EN^) (Fig. 4A-D). Live-cell STED imaging of CLCa and AP1µA revealed a perfect colocalization of AP-1 and clathrin on ARF1 compartments (Fig. 4B). In contrast, AP-3 and clathrin localized to segregated nanodomains (Fig. 4D). Colocalization analysis shows a strong overlap between AP-1 and clathrin, whereas AP-3 and clathrin colocalization was similar to the negative control (AP-1 vs. AP-3, Fig. 4E). To further elucidate the role of AP-1 on ARF1 compartments, we created a CRISPR-Cas9 AP1µA knock-out (KO) in the ARF1^EN^-Halo/SNAP-CLCa^EN^ KI cell line. Strikingly, loss of AP1µA led to the formation of elongated ARF1 compartments, suggesting a defect in fission (Fig. 4F). Clathrin recruitment to peripheral ARF1 compartments, but not to the Golgi, was impaired (Fig. 4G). Hence, AP-1 is responsible for the recruitment of clathrin and fission machinery to ARF1 compartments. Residual clathrin puncta were observed on ARF1 compartments (arrows in Fig. 4F) possibly suggesting recruitment via other clathrin adaptors such as GGAs (Golgi-localized, gamma-ear containing, ADP-ribosylation factor binding)^10^.

**Figure 4:**
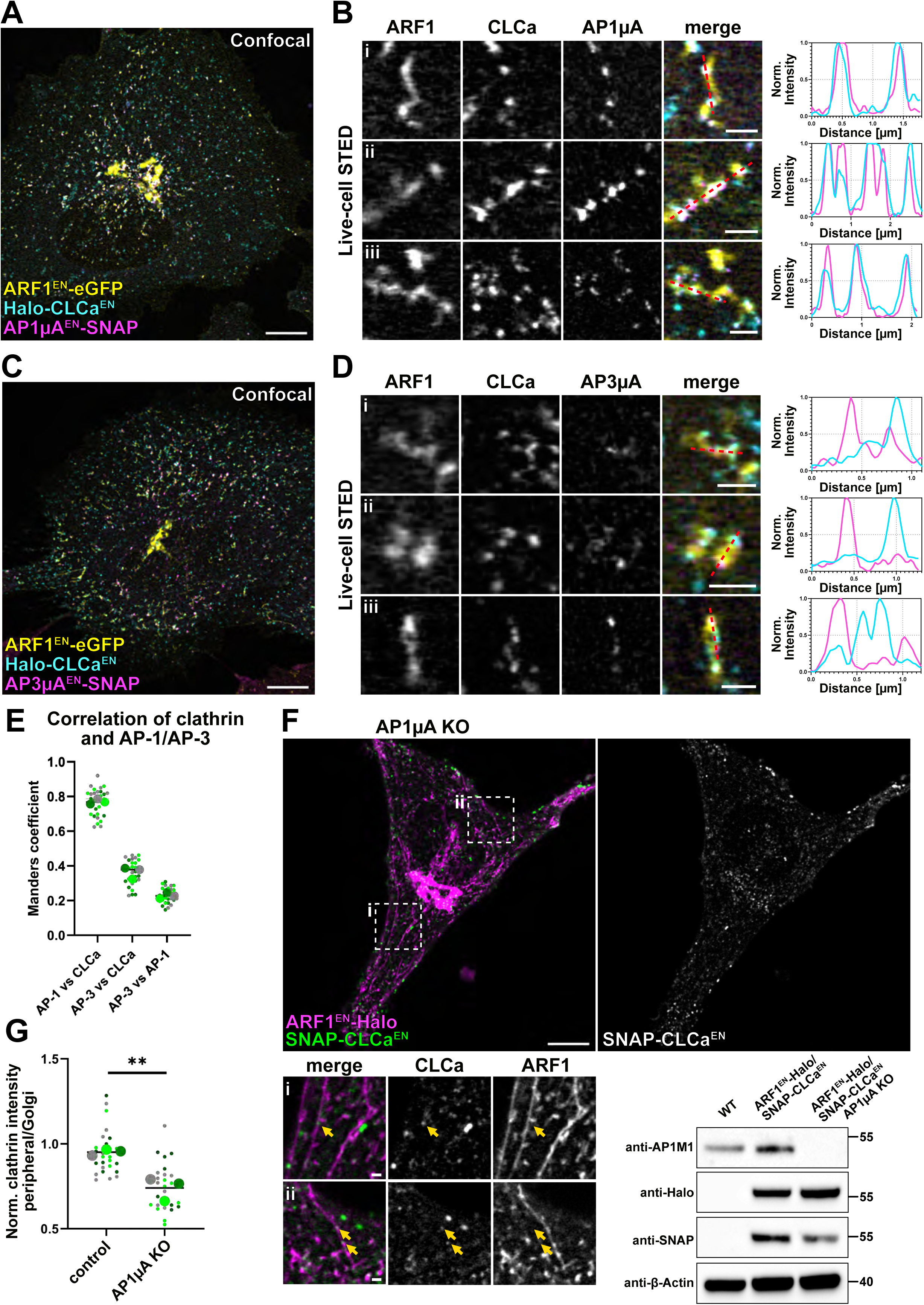
AP-1 recruits clathrin to ARF1 compartments and promotes their fission. (**A-B**) Live-cell confocal and STED imaging (2 color STED imaging with ARF1^EN^-eGFP imaged in confocal mode) of ARF1^EN^-eGFP/AP1µA^EN^-SNAP/Halo-CLCa^EN^ HeLa cells labeled with CA-JF_571_ and BG-JFX_650_ highlight that clathrin and AP-1 are recruited to the same nanodomains on ARF1 compartments. (i-iii) Examples of different compartments and line profiles showing perfect colocalization of AP-1 with clathrin. (**C-D**) The same analysis on ARF1^EN^-eGFP/AP3µA^EN^-SNAP/Halo-CLCa^EN^ HeLa cells labeled with CA-JF_571_ and BG-JFX_650_ shows that clathrin and AP-3 do not colocalize on ARF1 compartments. (i-iii) Examples of different compartments and line profiles. (**E**) Colocalization analysis using the Manders coefficient shows high correlation of AP-1 with clathrin but low correlation of AP-3 with clathrin, comparable with the correlation of AP-1 with AP-3. In total 30 cells from 3 independent experiments were analyzed, replicates are shown in different colors each dot representing a single cell. (**F**) Live-cell confocal imaging shows that AP1µA KO in ARF1^EN^-Halo/SNAP-CLCa^EN^ HeLa cells labeled with CA-JF_552_ and BG-JFX_650_ leads to the formation of aberrant long tubular ARF1 compartments. Clathrin recruitment to the long tubules is reduced but not completely abolished (yellow arrows highlight clathrin nano-domains in crops (i-ii). (**G**) Quantification of fluorescence intensity of clathrin punctae on peripheral ARF1 compartments normalized to the intensity of Golgi-associated clathrin punctae in control and AP1µA KO HeLa cells. In total 27 cells from 3 independent experiments were analyzed for each condition, replicates are shown in different colors and each dot represents a single cell of the replicate. P-value of nested t-test is 0.0064. EN=endogenous, BG=benzylguanine (SNAP-tag substrate), CA=chloroalkane (HaloTag substrate). Scale bars: 10 µm (confocal overview), 5 µm and 1 µm (crops).

To conclude, ARF1 compartments harbor AP-1 and AP-3 nanodomains, with clathrin being recruited exclusively to AP-1 nanodomains. The different classes of tubules, harboring different classes of adaptors may have a role in channeling different cargoes from ARF1 compartments into segregated downstream pathways. Beyond its role in cargo selection, AP-1 might also be required for the recruitment of fission factors.

### ARF1 compartments shed ARF1 to mature into Rab11-positive recycling endosomes

We found ARF1 compartments to harbor different AP complexes. As these adaptors were described to associate with different endosomal compartments^12,17,33^, we set out to investigate the nature of ARF1 compartments. For this, we visualized ARF1 in combination with known post-Golgi and endosomal membrane markers in live-cell confocal microscopy experiments (Fig. 5). First, we created a double KI cell line expressing edited ARF1 and Rab6, a membrane marker for tubular Golgi-derived carriers mediating direct transport to the plasma membrane^40^, and found ARF1 and Rab6 to define different post-Golgi carriers in the cell periphery (Fig. 5A). The high degree of colocalization at the Golgi prompted us to test the extent of overlap between Golgi-derived ARF1 compartments and Rab6 carriers (Extended Data Fig. 3). Live-cell STED and confocal time lapses show that about half of the Golgi-derived compartments are positive for ARF1 and Rab6, while the remaining half are Rab6-only carriers devoid of AP-1 (Extended Data Fig. 3A-C). This suggests the presence of functionally distinct classes of Rab6 carriers. Rab6-only tubules may be the direct Golgi-to-plasma membrane (PM) carriers, which have been reported previously^41^. Next, to test whether ARF1 compartments are defined by early, late, sorting or recycling endosomal markers, we created double KI cell lines expressing gene edited ARF1 and Rab7 (late endosome, LE), SNX1 (sorting endosome, SE) and Rab11 (recycling endosome, RE) or transiently overexpressed Rab5 (early endosome, EE). Live-cell confocal imaging revealed that ARF1 compartments are not defined by early, late or sorting endosomal markers, but they displayed a partial colocalization with the RE marker Rab11 (Fig. 5A-B). Interestingly, as reflected by the colocalization analysis, we observed ARF1 compartments positive for ARF1 only (Fig. 5Ai), RE structures positive for Rab11 only (Fig. 5Aii), and some ARF1 compartments which were also positive for Rab11 (Fig. 5Aiii). Accordingly, AP-1 and clathrin are seen on ARF1 compartments and on a sub-population of double-positive Rab11-ARF1 compartments (Extended data Fig. 4A-B). Fast confocal imaging showed co-translocation of AP-1 with ARF1 compartments only (Extended data Fig. 4C) and quantification of the overlap between ARF1/AP-1/Rab11 revealed a stronger association of AP-1 to ARF1 compartments compared to REs (Extended data Fig. 4D). In addition, we observed many ARF1 compartments that appear to localize closely to REs in confocal microscopy images (Fig. 5Aiv). To further characterize the dynamics of these various classes of ARF1 compartments and REs, we started out by surveying the dynamics of ARF1 compartments closely interacting with REs with higher resolution live-cell STED microscopy. Both peripheral and Golgi-derived compartments appeared to be closely associated and interacting with REs (Fig. 5C, Extended Data Fig. 5A-B, Extended Data Video 3-4). Furthermore, fast confocal microscopy and live-cell STED microscopy could highlight transient interaction of the same ARF1 compartment with different REs (Fig. 5D, Extended Data Fig. 5A-B). Interestingly, live-cell confocal microscopy revealed that AP-1 is located at the interface of ARF1 compartments and REs in cases where both compartments were found to interact (Extended Data Fig. 5C).

**Figure 5.**
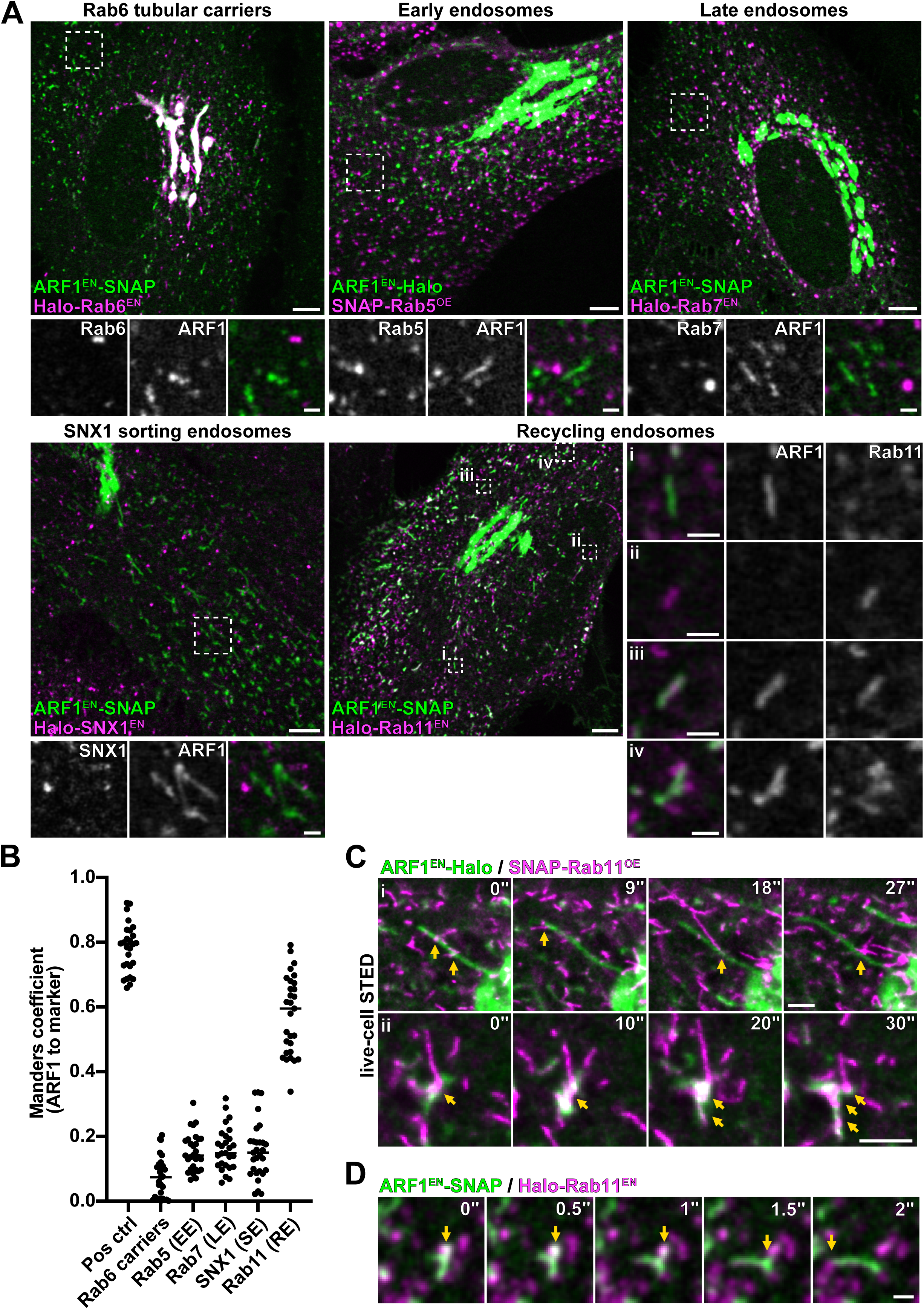
ARF1 compartments are novel sorting compartments with a partial colocalization with the RE marker Rab11. (**A**) Live-cell confocal microscopy of ARF1^EN^-SNAP/Halo-Rab6^EN^ (Rab6 secretory carriers), ARF1^EN^-SNAP/Halo-Rab7^EN^ (late endosomes), ARF1^EN^-SNAP/Halo-SNX1^EN^ (SNX1 sorting endosomes), ARF1^EN^-SNAP/Halo-Rab11^EN^ (recycling endosomes) HeLa cells labeled with BG-JF_552_ and CA-JFX_650_ (ARF1+Rab6/7/11) or BG-JFX_650_ and CA-JF_571_ (ARF1+SNX1). ARF1^EN^-Halo HeLa cells transiently expressing SNAP-Rab5^OE^ were labeled with BG-JFX_650_ and CA-JF_552_. Confocal imaging highlights ARF1 compartments devoid of markers for different endosomal compartments and a partial overlap of ARF1 compartments with the RE marker Rab11. In particular we could observe compartments positive for (i) ARF1 only (ii) for Rab11 only (iii) positive for both ARF1 and Rab11 (iv) and ARF1 compartments in close proximity to REs. **(B)** Colocalization analysis using the Manders coefficient shows higher correlation of ARF1 compartments with REs compared to other tested markers. At least 27 cells from 3 independent experiments were analyzed for each condition, each dot represents a single cell. (**C**) STED microscopy of ARF1^EN^-Halo HeLa cells transiently expressing SNAP-Rab11^OE^ labeled with CA-JF_571_ and BG-JFX_650_ show the dynamic nature of the interaction of ARF1 compartments with REs (sites of interaction indicated with yellow arrows) **(i)** at the TGN or **(ii)** in the cell periphery. (**D**) Time-lapse confocal spinning disk imaging in ARF1^EN^-SNAP/Halo-Rab11^EN^ HeLa cells labeled with CA-JFX_650_ and BG-JF_552_ shows ARF1 compartments transiently interacting with different REs (sites of interaction indicated with yellow arrows). EN=endogenous, BG=benzylguanine (SNAP-tag substrate), CA=chloroalkane (HaloTag substrate). Scale bars: 5 µm (overview) and 1 µm (crops, time-lapse).

Yet, we were intrigued by the fact that some ARF1 compartments were also defined by Rab11 (Fig. 5Aiii), raising the possibility that ARF1 compartments may mature into REs. To test this, we endogenously tagged ARF1 with the highly photostable monomeric fluorescent protein StayGold (mStayGold)^42^, allowing long term imaging of ARF1 and Halo-Rab11 dynamics (Fig. 6). We focused on peripheral ARF1 compartments as the cell periphery is a less crowded environment allowing to follow structures over time. Strikingly, we observed that double positive compartments shed their ARF1 coat and acquired more Rab11 over time (Fig. 6A-C, Extended Data Video 5). We observed ∼ 1-2 maturation events per ∼ 200 µm^2^ of cell area in ∼ 4 minutes acquisition time. While increase of Rab11 coating the surface of the compartment was gradual, complete shedding of ARF1 occurred over a short period of ∼7-9 seconds (Fig 6B). Noticeably, the highly curved ends of the ARF1 compartment were the last parts to uncoat (Fig. 6B yellow arrows), suggesting that ARF1 molecules may be shielded by the presence of the AP-1/clathrin coat which was observed to preferentially localize to regions of high curvature on ARF1 compartments. Taken together, these data suggest that ARF1 compartments mature into REs to possibly be able to subsequently deliver their content to the PM.

**Figure 6.**
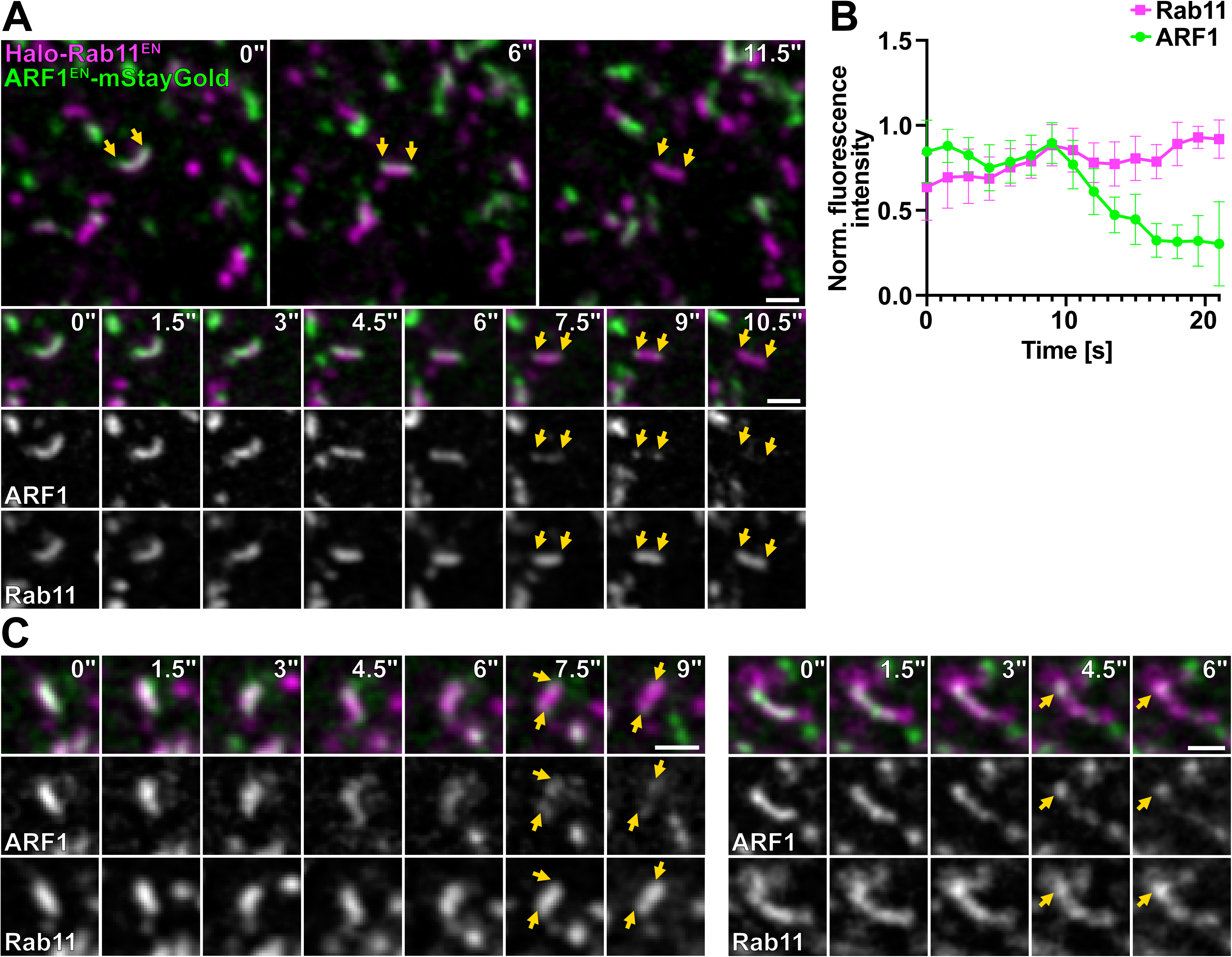
ARF1 compartments mature into Rab11-positive REs. (**A**) Live-cell confocal microscopy of ARF1^EN^-mStayGold/Halo-Rab11^EN^ HeLa cells labeled with CA-JFX_650_. ARF1 compartments are seen to shed ARF1 from their membrane and mature into REs. (**B**) Normalized fluorescence intensity of ARF1 and Rab11 signal on maturing endosomal compartments. ARF1 is shed over a short period of 7-9 s. Error bars display SD. (**C**) Additional examples of maturation events of ARF1 compartments into REs. Same conditions as in **A**. Yellow arrows indicate the curved ends of the tubules which uncoat last. EN=endogenous, CA=chloroalkane (HaloTag substrate). Scale bars: 1 µm (crops, time-lapse).

### ARF1 compartments mediate endocytic recycling and secretory traffic via maturing into recycling endosomes

So far, we have shown that ARF1-positive compartments are novel organelles that control clathrin and adaptor-dependent post-Golgi trafficking. To determine the exact role of ARF1 compartments in sorting, we investigated the trafficking of endocytic and secretory cargoes in different CRISPR-Cas9 KI cell lines. We first started by investigating the role of Golgi-derived perinuclear ARF1 compartments in the export of cargoes from the Golgi (Fig. 7). ARF1 compartments have been shown to mediate Golgi export of vesicular stomatitis virus glycoprotein (VSV-G)^13^. Interestingly, they were not observed fusing with the PM suggesting that another sorting step is required for the final delivery of VSV-G to the PM. We employed the Retention Using Selective Hooks (RUSH) system to release a pulse of various secretory cargoes from the ER^43^. The RUSH system consists of a reporter fused to a fluorescent protein or SNAP-tag and to streptavidin binding protein (SBP). The reporter is retained in the ER by a streptavidin hook. Upon biotin addition, the reporter is released from the ER and accumulates at the Golgi (∼15 min) before reaching the PM (∼30 min). We examined the secretory transport of five different RUSH reporters: transferrin receptor (TfR, transmembrane protein), a LAMP1 variant lacking the endocytic motif in its cytoplasmic tail and thus fails to be endocytosed after deposition to the PM (LAMP1△, transmembrane protein)^44^, TNFα (transmembrane protein), VSV-G (transmembrane protein) and a soluble SNAP reporter (sSNAP) with live-cell confocal microscopy in the various gene-edited cell lines. We found that all reporter cargoes exited the Golgi in ARF1/clathrin compartments that did not harbor AP-3 (Fig. 7A-B, Extended Data Fig. 6A-C, Extended Data Video 6). Secretory compartments detached and moved away from the Golgi (Extended Data Fig. 6D). Using two exemplary RUSH cargoes, we then quantified the fraction of the tubulo-vesicular carriers leaving the Golgi that were positive for ARF1. We found that ∼ 90% of RUSH cargo-containing tubules were decorated by ARF1 (Fig. 7C).

**Figure 7.**
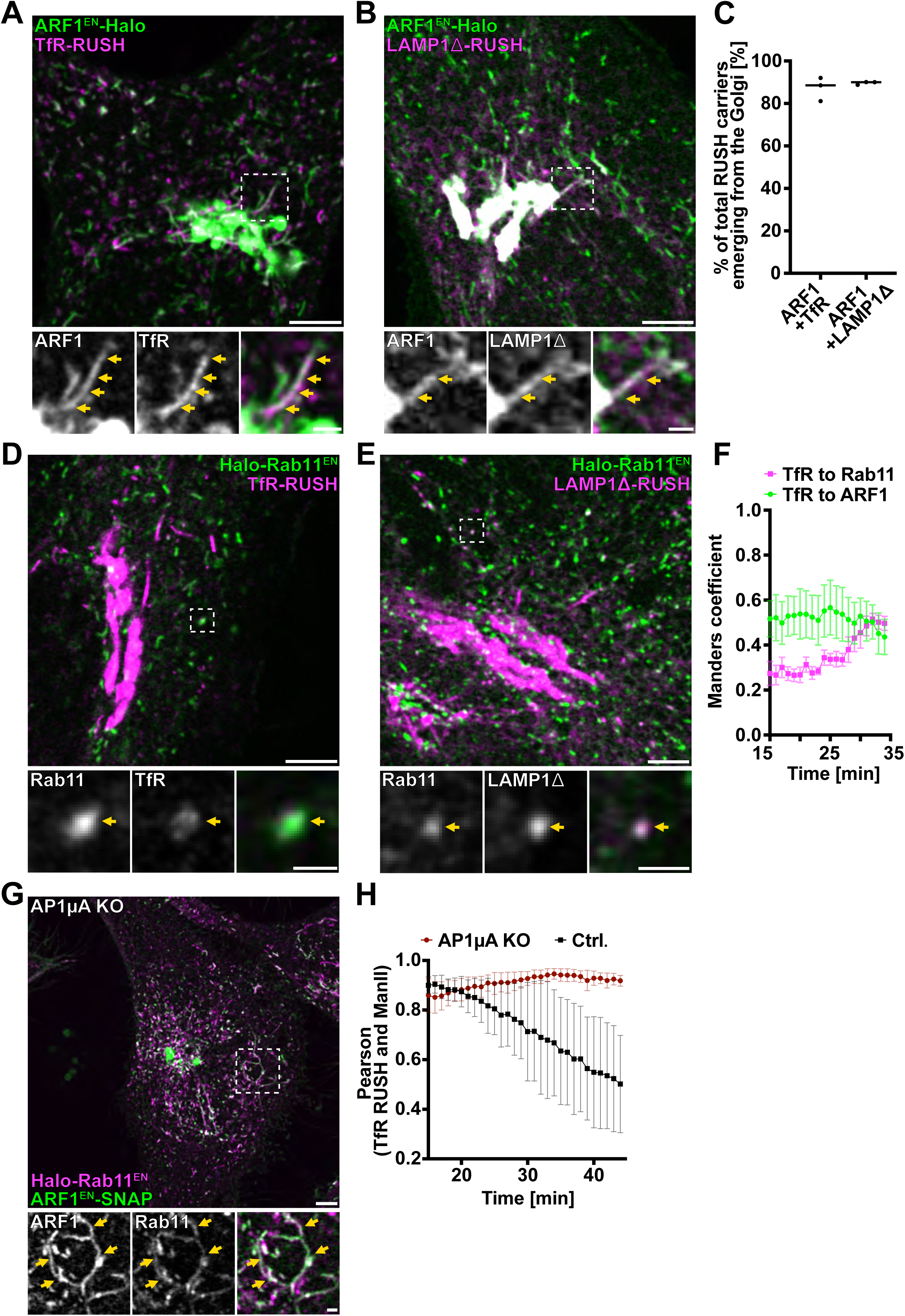
Perinuclear ARF1 compartments transport secretory cargoes and loss of AP-1 delays cargo exit from the TGN. (**A-B**) ARF1^EN^-Halo HeLa cells transiently expressing the RUSH constructs (**A**) Streptavidin-KDEL/TfR-SBP-SNAP or (**B**) Streptavidin-KDEL/ssSBP-SNAP-LAMP1Δ (w/o QYTI) labeled with CA-JF_552_ and BG-JFX_650_ were imaged with confocal microscopy 20 minutes after addition of biotin with a frame rate of 6 s/frame. A qualitative example shows secretory RUSH cargo leaving the Golgi in ARF1 compartments at 21 min (TfR) or 24 min (LAMP1ϕλ) post biotin. (**C**) Manual quantification of the total RUSH cargo carriers emerging from the Golgi reveals that most secretory RUSH cargo exits via ARF1 compartments, each dot represents a single cell. (**D-E**) Live-cell confocal imaging of Halo-Rab11^EN^ HeLa cells transiently expressing (**D**) Streptavidin-KDEL/TfR-SBP-GFP labeled with CA-JFX_650_ and (**E**) Streptavidin-KDEL/ssSBP-SNAP-LAMP1Δ labeled with CA-JF_552_ and BG-JFXX_650_ shows that secretory RUSH cargo is sorted through REs *en route* to the PM. A qualitative example is at 21 min (TfR) or 24 min (LAMP1ϕλ) post biotin (**F**) Colocalization analysis using the Manders coefficient shows that correlation of TfR-RUSH cargo with REs increases over time while correlation with ARF1 compartments shows a downward trend. Each dot represents the average of 5 cells, SEM error bars. (**G**) Live-cell confocal imaging of ARF1^EN^-SNAP/Halo-Rab11^EN^ AP1µA KO HeLa cells labeled with CA-JF_552_ and BG-JFX_650_, exhibit formation of long aberrant Rab11/ARF1-compartments near the TGN. (**H**) Pearson correlation coefficient of secretory TfR-RUSH cargo and the Golgi (masked by the Golgi-marker ManII) reveals that upon AP1µA KO, Golgi exit of secretory cargo is impaired in comparison to control cells, each dot represents the average of 4 cells, SD error bars. EN=endogenous, BG=benzylguanine (SNAP-tag substrate), CA=chloroalkane (HaloTag substrate). Scale bars: 5 µm (overviews) and 1 µm (crops).

REs have been shown to serve as an intermediate sorting station for secretory cargo exiting the Golgi *en route* to the PM^45,46^. Since ARF1 compartments do not fuse with the PM^13^ but shed ARF1 to mature into REs (Fig. 6), we tested whether secretory RUSH cargo transit through REs downstream of ARF1 compartments. For this, we followed the transport of RUSH cargoes in Halo-Rab11^EN^ KI cells and found them to colocalize with REs (Fig. 7D-E and Extended Data Fig. 6E). By quantifying the kinetics of TfR RUSH transport to ARF1 compartments and REs, we found that the cargo first fills ARF1 compartments before transitioning to REs (Fig. 7F). These results suggest a model in which ARF1 compartments containing secretory cargo emerge from the Golgi and over time mature into REs before being able to deliver their content to the PM. Furthermore, upon AP1µA KO, both TfR- and TNFα-RUSH were retained in long aberrant perinuclear tubules that were positive for both ARF1 and Rab11 (Fig. 7G and Extended Data Fig. 6F). Additionally, cargo exit from the TGN was delayed in AP1µA KO cells (Fig. 7H). Altogether, this suggests that AP-1 may be required for secretory ARF1 compartments to shed ARF1 and mature into REs.

As secretory cargoes were almost exclusively transported by perinuclear ARF1 compartments, we wondered about the function of peripheral ARF1 compartments. AP-1 was previously shown to be important for transferrin (Tfn) recycling^47^, motivating us to investigate the role of ARF1 compartments in endocytic recycling (Fig. 8). For this, we performed fluorescent Tfn uptake experiments in different gene-edited cell lines. After internalization, Tfn could be detected in peripheral ARF1 compartments (Fig. 8A) defined by AP-1 and AP-3 nanodomains (Extended Data Fig. 7A-C). These peripheral ARF1 compartments were functionally segregated from the perinuclear secretory ARF1 compartments (Fig. 8B). Quantification of the colocalization of fluorescent Tfn with markers for EE (Rab5), ARF1 compartments and REs (Rab11) showed that internalized Tfn first localizes to Rab5-positive EEs, then to ARF1 compartments and REs (Fig. 8C). 10 mins post-internalization Tfn empties out from ARF1 compartments but continues to fill REs (Fig. 8C). Dynamic live-cell microscopy further demonstrates that ARF1 compartments are not simply sorting sub-domains of EEs or derive from EEs (Extended Data Fig. 7D).

**Figure 8.**
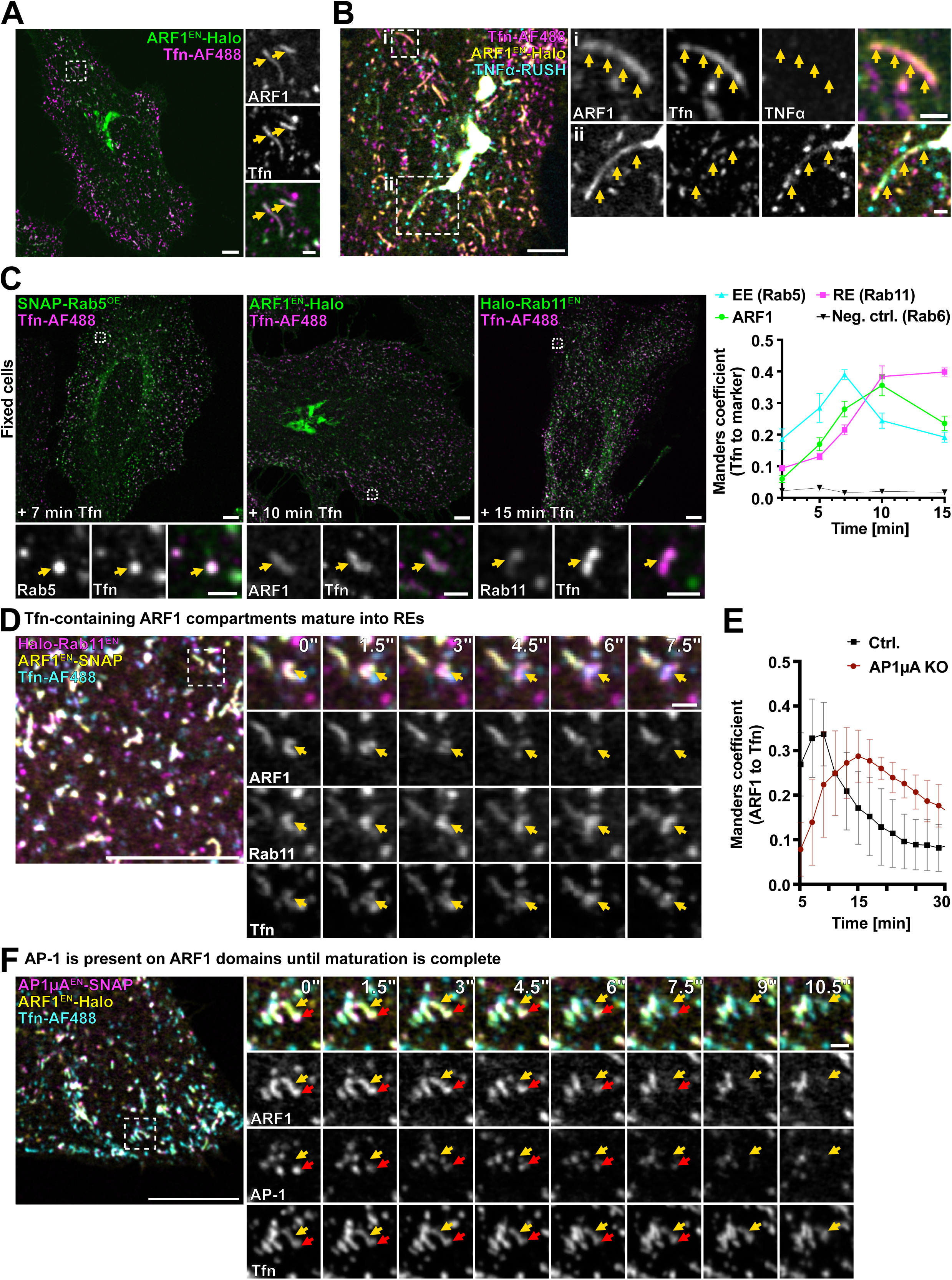
ARF1 compartments mediate endocytic recycling and direct cargo flow via maturation into REs. (**A**) Transferrin (Tfn) recycling assays were performed using fluorescently labeled Tfn (Tfn-AlexaFluor488). Live-cell confocal imaging in ARF1^EN^-Halo HeLa cells labeled with CA-JFX_650_ shows Tfn in ARF1 compartments 5 min after addition of Tfn. (**B**) Live-cell confocal imaging of ARF1^EN^-Halo HeLa cells transiently expressing Streptavidin-KDEL/TNFα-SBP-SNAP labeled with BG-JFX_650_ and CA-JF_552_ shows that when performing both RUSH and Tfn recycling assay in parallel, both cargoes are in separate ARF1 compartments: (i) Peripheral ARF1 compartments containing only endocytic recycling cargo and (ii) perinuclear ARF1 compartments containing only secretory RUSH cargo. (**C**) Tfn recycling assays using Tfn-AlexaFluor488 were performed in ARF1^EN^-Halo, Halo-Rab6^EN^, Halo-Rab11^EN^ HeLa cells and Hela cells transiently expressing SNAP-Rab5 (SNAP-Rab5^OE^) labeled with CA-JFX_650_ or BG-JFX_650_. Cells were fixed at indicated timepoints post addition of Tfn to the culture media. Colocalization analysis using the Manders correlation coefficient showed that Tfn first enters EE, then ARF1 compartments and REs, Halo-Rab6^EN^ cells were used as a negative control. Each data point represents the average of 10 cells of two independent experiments, SEM error bars. (**D**) Live-cell confocal microscopy of Tfn recycling using Tfn-AlexaFluor488 in ARF1^EN^-SNAP/Halo-Rab11^EN^ HeLa cells labeled with BG-JFX_552_ and CA-JFX_650_. Tfn-containing ARF1 compartments are seen to shed ARF1 from their membrane and mature into REs. (**E**) Correlation analysis using the Manders coefficient of ARF1 and Tfn in ARF1^EN^-Halo HeLa cells or ARF1^EN^-Halo/AP1µA KO HeLa cells shows that Tfn retains longer in ARF1 compartments upon KO of AP1µA. Each dot represents the average of 5 cells, SD error bars. (**F**) Live-cell confocal microscopy of Tfn recycling using Tfn-AlexaFluor488 in ARF1^EN^-Halo/AP1µA^EN^-SNAP HeLa cells labeled with BG-JFX_552_ and CA-JFX_650_. AP-1 localizes to maturing ARF1 compartments which contain Tfn. EN=endogenous, BG=benzylguanine (SNAP-tag substrate), CA=chloroalkane (HaloTag substrate). Scale bars: 5 µm (overviews **A, B, C**), 10 µm (overviews in **D** and **F**) and 1 µm (crops).

We then wanted to test whether flow of Tfn cargo is dependent on the maturation of ARF1 compartments into REs. For this, we applied fluorescent Tfn to Rab11 and ARF1 double KI cells and followed the fate of an ARF1/Rab11 compartment filled with Tfn. Over the time of ∼ 9 s ARF1 dissociated from the membrane of the Tfn filled compartment resulting in the formation of a Rab11-only positive RE (Fig. 8D, Extended Data Video 7). Loss of AP1µA delayed Tfn export from ARF1 compartments (Fig. 8E). Intriguingly, ARF1 compartments involved in Tfn recycling did not appear elongated as a result of the AP1µA KO (Extended Data Fig. 7E), suggesting that fission defects caused by the loss of AP-1 are limited to perinuclear secretory ARF1 compartments. Furthermore, we observed that AP-1 stays associated with ARF1 compartments during the maturation process and dissociates from the membrane once maturation is complete (Fig. 8F, Extended Data Video 8).

Collectively, these data suggest that ARF1 compartments mediate endocytic recycling via maturation into REs, a process that depends on AP-1.

## Discussion

Clathrin-coated vesicles are thought to mediate cargo exchange between the TGN and different endosomal compartments as small punctate structures are commonly seen travelling around the cytoplasm of cells^6–8^. Endogenous tagging of ARF1 and clathrin together with CLEM allowed us to visualize the membrane underlying clathrin-positive vesicle-like structures and revealed that the vast majority of non-endocytic clathrin is associated with a tubulo-vesicular network of ARF1 compartments (Fig. 1-2) which direct cargo flow along the secretory and endocytic recycling routes though shedding of ARF1 and maturation into recycling endosomes (Fig. 9). Previous studies have shown clathrin as well as AP-1 and AP-3 on tubular endosomal structures, however the identity and the role of these compartments was never assessed further^14,16,17^. Here we report functionally different classes of ARF1 compartments defined by different combinations of AP nanodomains: perinuclear ARF1 compartments with a secretory function that harbor AP-1 only and peripheral endocytic recycling compartments that harbor both AP-1 and AP-3 (Fig. 3). Clathrin translocates and remains associated with ARF1 compartments, and no budding events are observed (Fig. 1). However, visualization of budding events from moving objects is technically challenging and would require fast volumetric imaging to detect rapid uncoating events after vesicle formation.

**Figure 9.**
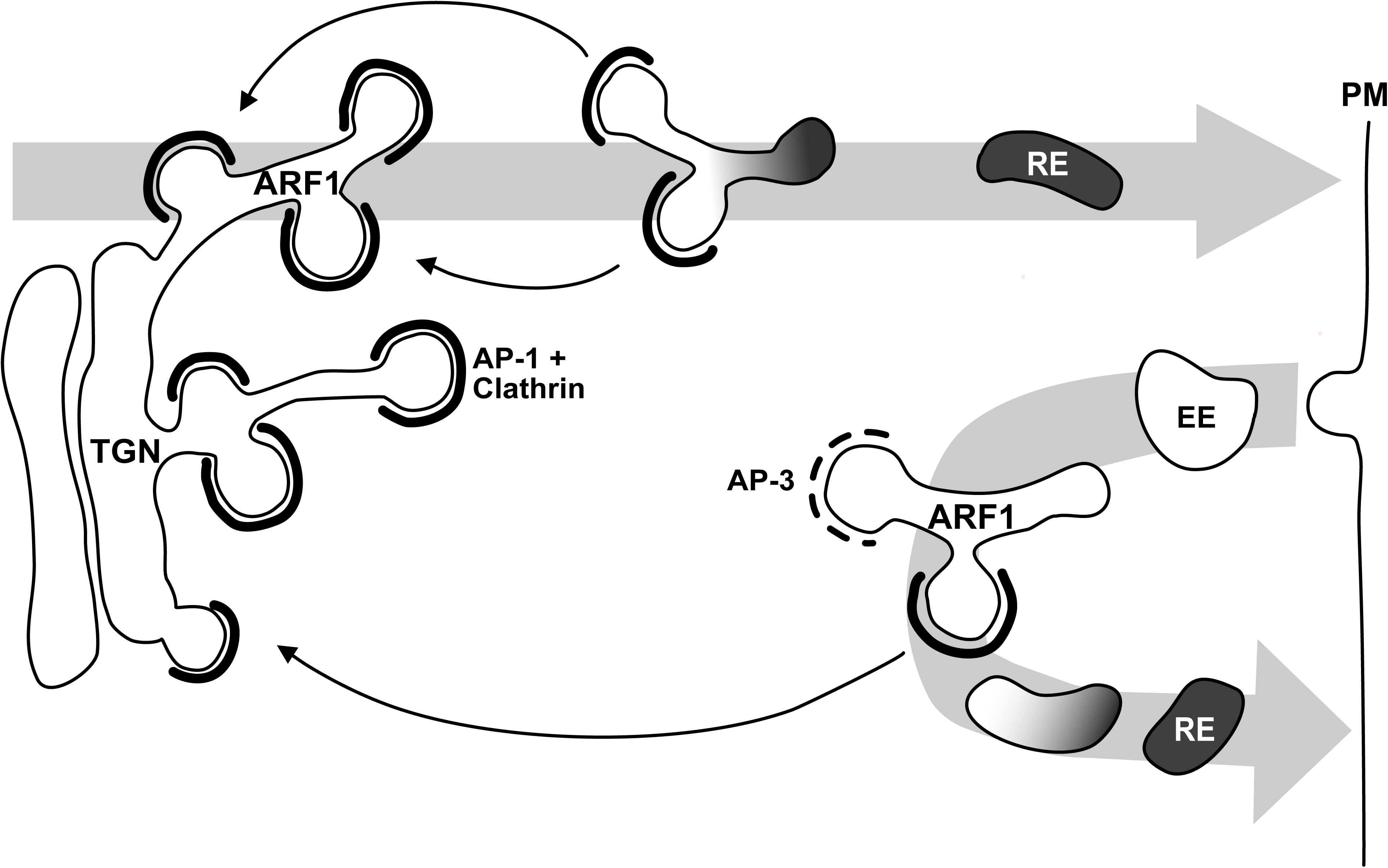
ARF1 compartments orchestrate cargo flow via maturing into RE. Clathrin-dependent post Golgi pathways are mediated by different ARF1 compartments. Secretory and endocytic recycling cargoes flow would be driven by maturation of ARF1 compartments into REs, while endosome-to-Golgi transport would be driven by AP-1 carriers. Functionally different classes of ARF1 compartments harbor AP nanodomains allowing for site-specific cargo enrichment. RE = recycling endosome EE = early endosome, TGN = trans-Golgi network. We could observe tubulo-vesicular ARF1 compartments decorated with clathrin in different mammalian cell types (Extended Data Fig. 1), hinting at a conserved mechanism for post-Golgi transport via ARF1 compartments. The concept of endosomal maturation has already been described, as early endosomes were seen to shed Rab5 and acquire the late endosome marker Rab7^21,22^. We here propose a similar mechanism for conversion of ARF1 compartments into recycling endosomes, which may be driven by an ARF-Rab cascade as shown for the Rab5 to Rab7 conversion. It remains elusive whether ARF1-to-Rab11 conversion is important for the biogenesis of all recycling endosomes. We envision that the membranes may emerge from the ER-Golgi interface and generate ARF1 compartments via complex fission and fusion mechanisms that may require fast volumetric imaging to postulate a compelling model. Moreover, the mechanisms of cargo transfer from EE to ARF1 compartments need to be explored, as our data suggest that ARF1 compartments do not directly bud from EEs (Extended Data Fig. 7D).

Overall ARF1 compartments are not defined by endosomal markers, but partial colocalization was observed with Rab11, a Rab GTPase commonly used as a RE marker (Fig. 5A-B). We propose that ARF1 compartments represent a novel tubolo-vesicular organelle with a key role in the distribution of secretory and endocytic recycling cargo along post-Golgi routes. While vesicular exchange between the Golgi and endosomes has been proposed, our data rather suggest that maturation of ARF1 compartments into REs directs the flow of anterograde cargoes exiting the Golgi (Fig. 6-8). Our findings close an important gap in the understanding of communication between the Golgi and post-Golgi organelles. Peripheral ARF1 compartments are the sorting station downstream of Rab5 EEs (Fig. 8C). Dynamic imaging shows that ARF1 compartments are not simply tubulo-vesicular compartments derived from EEs but rather a stand-alone organelle (Extended Data Fig. 7D). Peripheral ARF1 compartments containing fluorescent transferrin are seen shedding ARF1 and acquiring the identity of REs, explaining that re-deposition of cargoes back to the PM is mediated by maturation (Fig. 8D). It is likely that ARF1 compartments emerging from the Golgi (which are also marked by Rab6) would undergo a similar transition to acquire endosomal identity and would shed ARF1 before being able to fuse with the PM. Previous reports in yeast and *Drosophila melanogaster* have postulated the presence of a Rab6-to-Rab11 cascade at the Golgi and for dense granule and exosome biogenesis suggesting this GTPase switch may be conserved across kingdoms^48,49^. Some secretory cargoes have been shown to follow an indirect route from the Golgi to the plasma membrane via REs^45,46^ and the maturation of TGN-derived ARF1 compartments into Rab11-positive endosomes explains why cargoes are observed in endosomes downstream of the Golgi. Unfortunately, the detection of maturation events of TGN-derived tubules filled with secretory cargoes is challenging since these tubules move in and out on the plane on their way from the Golgi to the PM. However, the comparable behavior of the compartments would suggest similar sorting mechanisms. Additionally, a maturation defect could explain the formation of long-aberrant Rab11 and ARF1 positive tubules filled with secretory cargo upon KO of AP-1 (Fig. 7G, Extended Data Fig. 6F). Impaired TGN export could additionally be explained by a defect in the retention of TGN-resident proteins or possibly defective retrograde endosome-to-Golgi recycling^34^.

What is the role of AP-1 on ARF1 compartments? AP-1 is a versatile adaptor that acts in many trafficking steps. Because of this plethora of functions in many different cell types, it has been difficult to reach a consensus about the core functions of AP-1. AP-1 is thought to coordinate bi-directional transport between the TGN and endosomes^1,33,34^. In yeast, an additional role for AP-1 in intra-Golgi recycling of Golgi-resident proteins has also been suggested^50^. Recently, AP-1 was proposed to act exclusively in retrograde transport of cargoes from endosomes back to the Golgi^14^. Our data show that AP-1 localizes solely on the Golgi and ARF1 compartments. Assuming that the role of AP-1 is connected to protein transport back to the Golgi apparatus, it is conceivable that a compartment may have to retrieve all retrograde cargoes before becoming competent for transport to the PM (Fig. 9). AP-1 could sequester AP-1 cargoes from ARF1 compartments, whether they have escaped the Golgi (perinuclear compartments) or upon internalization after endocytosis (peripheral compartments). Once retrograde cargoes are exported from ARF1 compartments, ARF1 would dissociate from the membrane. Interestingly, shedding of ARF1 from the membrane appears to be slowest at the regions of high curvature on the compartment where AP-1/clathrin localize (Fig. 6 and 8F), suggesting that the coat may shield ARF1 from GTP hydrolysis. We speculate that ARF1 shedding would be triggered by recruitment of specific ARF-GAPs, possibly recruited by Rab11. Interestingly, ARF1 compartments are seen closely interacting with Rab11-positive recycling endosomes (Fig. 5 and Extended Data Fig. 5) with a similar behavior to that observed for other endosomal compartments that undergo kiss-and-run for cargo exchange^9,28^. We speculate that such close interactions may be necessary for ARF1 compartments to acquire maturation factors and/or allow material exchange. However, further experiments would be required to prove this concept.

The presence of AP-1 and AP-3 nanodomains on ARF1 compartments involved in Tfn recycling (Extended Data Fig. 7A-C) tempts us to speculate that AP-3 could, in a similar manner, direct transport of endocytosed proteins to endo-lysosomes from peripheral ARF1 compartments. AP-3 is reported to act in sorting from endosomes to melanosomes but its role in non-specialized cells is controversial^32^. Generally, AP-3 might be involved in the sorting of lysosomal cargoes as LAMP1 is mistargeted in AP-3 deficient mice^51^, calling for further investigation on the role of ARF1 compartments in transport to lysosomes. Although it was speculated that AP-3 does not bind clathrin *in vivo*^37,38^, the presence of a clathrin box in the β-subunit of AP-3 as well as its ability to bind clathrin *in vitro* indicated a possible AP-3/clathrin interaction^6^. Interestingly, we only see association of clathrin with AP-1 but not with AP-3 (Fig. 4A-E), indicating that AP-3 and clathrin do not interact *in vivo*.

Upon AP1µA KO we observed aberrant secretory ARF1 compartment containing anterograde cargo suggesting a fission defect (Fig. 7G-H, Extended Data Fig. 6F). We speculate AP-1 is responsible for the recruitment of yet-to-be identified fission factors. A potential role for the membrane-remodeling GTPase dynamin in fission events at the TGN is controversial, as several approaches suggested a contribution of dynamin to the fission of clathrin-coated vesicles from the TGN and endosomes^52–54^, while *in vivo* studies show that it exclusively promotes fission of endocytic vesicles^55,56^. Interestingly, ARF1 has been proposed to mediate dynamin 2 recruitment, and depletion of dynamin 2 led to long tubular extensions from the TGN^53^, reminiscent of the elongated ARF1 compartments we observe upon AP-1 depletion. Notably, peripheral ARF1 compartments involved in Tfn recycling remain morphologically unchanged upon AP1µA KO, while Tfn recycling is affected (Fig. 8E, Extended Data Fig. 7)^47^. Differential recruitment and specific function of AP-1 may be driven by distinct subsets of interacting proteins and co-adaptors, including specific lipid-modifying enzymes such as phosphatidylinositol-4-kinases^57^. Loss of AP1µA led to a pronounced reduction of clathrin association on long aberrant ARF1 compartments, without affecting clathrin recruitment to the Golgi (Fig. 4F-G). Clathrin could be recruited to ARF1 compartments or the TGN via other ARF-dependent adaptors such as GGA proteins^34^. Sub-populations of AP-1, either associated with GGAs or alone could drive differential effector recruitment. Additionally, the role of non-classic adaptor proteins, such as EpsinR^58^, in intracellular clathrin-dependent sorting is not well explored.

As the organization of the endosomal system differs strongly between organisms and cell types, due to the presence of specialized endosomes and distinct metabolic needs, we expect cell type-dependent variations of the here described mechanism^59^. ARFs are abundant and ubiquitously expressed proteins in all organisms, suggesting their key role in post-Golgi trafficking may be conserved. Diversity of function in different cell types and organisms might also be driven by the presence of different adaptors and effector proteins. In polarized epithelia cells, the tissue-specific adaptor AP-1B is additionally expressed^60^, adding another layer of complexity to the sorting process.

Overall, by using advanced imaging methods to visualize endogenously labeled sorting machinery, we provide evidence for intracellular material exchange being facilitated by a tubulo-vesicular network that connects the TGN, the endosomal system and the PM and drives cargo flow via maturation.

## Methods

### Mammalian cell culture

CCL-2 HeLa cells (ECACC General Collection) and HAP1 cells^61^ were grown in a humidified incubator at 37°C with 5% CO_2_ in Dulbecco’s Modified Eagle Medium (DMEM, Gibco) supplemented with 10% fetal bovine serum (Corning), 100 U/L penicillin and 0.1 g/L streptomycin (FisherScientific). Jurkat T-cells (DSMZ) were grown in a humidified incubator at 37°C with 5% CO_2_ in Roswell Park Memorial Institute medium (RPMI, Gibco) supplemented with 10% fetal bovine serum (Corning), 100 U/L penicillin and 0.1 g/L streptomycin (FisherScientific).

For transient transfection of plasmids encoding for SNAP-Rab5, GFP-ManII or RUSH cargoes into HeLa cells, a NEPA21 electroporation system was used (Nepa Gene). For this, 1 million cells were washed twice with Opti-MEM (Gibco), resuspended in 90 µl Opti-MEM and mixed with 5 µg for SNAP-Rab5, 1 µg for ManII-GFP, 10 µg for RUSH cargo of DNA in an electroporation cuvette with a 2-mm gap. The electroporation reaction consists of two poring pulse (125 V, 3 ms length, 50 ms interval, with decay rate of 10% and + polarity) and five consecutive transfer pulse (25 V, 50 ms length, 50 ms interval, with a decay rate of 40% and ± polarity).

### Recombinant plasmids for overexpression

See supplementary information.

### CRISPR-Cas9 KI and KO plasmids and cell lines

See supplementary information.

### Live-cell labeling

For live-cell imaging, cells were seeded on a glass-bottom dish (3.5 cm, #1.5; Cellvis) coated with fibronectin (Sigma). Labeling with Halo and SNAP substrates was carried out for 1 h at 37°C in culture media. All dyes-conjugates were used at concentration of 1 µM. After the staining, cells were washed in growth media at 37 °C for at least 1 h. Live-cell imaging was performed in live-cell imaging solution (FluoBrite DMEM (Gibco) supplemented with 10% FBS, 20 mM HEPES (Gibco) and 1x GlutaMAX (Gibco)).

For live-cell labeling of non-adherent T-cells, 200 000 cells were labeled with Halo and SNAP substrates (1µM) for 1 h at 37°C in culture media in a volume of 100µl. After staining, cells were washed in growth media 3 times via centrifugation.

### Imaging and image processing

Line-scanning confocal and STED imaging was carried out on a commercial expert line Abberior STED microscope equipped with 485 nm, 561 nm and 640 nm excitation lasers. For two-color STED experiments both dyes were depleted with a 775 nm depletion laser. The detection windows were set to 498 to 551 nm, 571 to 630 nm and 650 to 756 nm. Multi-color STED images were recorded sequentially line by line. For confocal imaging, probes that were detected in the 498 to 551 nm and the 650 to 756 nm detection windows were recorded simultaneously. If required for quantitative analysis, laser power was kept constant between images. For live-cell confocal imaging the pixel size was set to 60 nm and for STED imaging to 30 nm (live-cell) or 20 nm (fixed cell). Live-cell imaging was performed at 37°C.

Spinning disk confocal imaging was carried out at CSU-W1 SoRa spinning disk (Nikon) with a dual camera system for simultaneous dual-color detection. All imaging was performed with a 60x Plan Apo oil objective (NA=1.4). For experiments 488 nm, 561 nm and 636 nm laser lines were used for excitation. For simultaneous dual color imaging, the quad-bandpass filter was used with relevant center wavelengths of 607 nm and 700 nm and full-width half maximums of 34 nm and 45 nm, respectively. For three-color imaging, eGFP and far-red (JFX_650_) signal were detected simultaneously, and the orange channel (JF_552_) was detected separately. Again, the quad-bandpass filter was used with relevant center wavelengths of 521 nm, 607 nm and 700 nm and full-width half maximums of 21 nm, 34 nm and 45 nm, respectively.

To reduce noise, confocal images were background subtracted and gaussian blurred (1 pixel SD) using Fiji^62^. STED videos in Fig. 5C and images in Extended Data Fig. 3A, 4A-B, 5A-B were deconvoluted using Richardson–Lucy deconvolution from the python microscopy PYME package (https://python-microscopy.org). Line profiles shown in Fig. 4B, D and Extended Data Fig. 5A were obtained by drawing a perpendicular line to the direction of the membrane with Fiji on gaussian blurred (1 pixel SD) STED images. The line profile data was then normalized and plotted using GraphPad Prism (GraphPad Software, https://www.graphpad. com).

## FIB-SEM CLEM

For 3D CLEM, ARF1^EN^-Halo/SNAP-CLCa^EN^ HeLa cells were grown on IBIDI gridded glass coverslips. Cells were stained with SNAP-tag and HaloTag substrates as for standard light microscopy. Additionally, lysosomes were labeled SiX-lysosome^63^ (0.5 µM) after Halo- and SNAP-labeling was complete, to visualize lysosomes. Cells were fixed for 15 min at 37°C using 3% PFA and 0.2% glutaraldehyde. Following the fixation, the reaction was quenched using 0.1% NaBH_4_ in PBS for 7 min and cells were rinsed three times with PBS afterwards. Cells were then directly imaged in PBS. From chosen cells confocal z-stacks with 200 nm increments were recorded. All three colors were recorded sequentially line-by-line.

Following light microscopy, cells were postfixed with 2% glutaraldehyde in 0.1 M cacodylate buffer. Osmification was performed with 1% OsO4 and 1.5% potassium cyanoferrate (III) in 0.1 M cacodylate buffer on ice, followed by aqueous 1% OsO4 at room temperature. Following washing, 1% uranyl acetate was applied and dehydration in ascending ethanol concentration was performed. Ultrathin embedding into Durcupan resin was performed^64^. After centrifugation of excessive resin amount, coverslips with cells were put into the heating cupboard for polymerization.

Following resin polymerization, coverslips were trimmed with a glass cutting pen and mounted onto SEM pin stabs, sputter coated with carbon (30-50 nm) and imaged in SEM (Helios 5CX). Low-resolution overviews of coverslip surface were used to navigate on the coverslip and find the corresponding quadrant with the cell of interest. Sum projection image from SDCM was overlaid with a SEM cell image (ETD, secondary electrons) in Thermo Fischer MAPS software to specify part of cell for autoslice and view (ASV). FIB milling was performed at 7 nm step, SEM imaging was performed at x = 3.37 nm; y = 4.6 nm, dwelling time 5 µs, backscattered electrons were detected by in-Column detector at 2kV, 0.34 nA.

Stack alignment was performed in freeware Microscopy image browser (MIB). 3D stacks were binned to 7 nm isotropic resolution and further CLEM alignment, feature segmentation and visualization was performed using Dragonfly software. In short, SiX-labeled lysosomes were used for rough alignment, fine alignment of light microscopy and electron microscopy volumes was performed by orienting on cell membrane edges and plasma membrane coated pits, that were clearly visible in FIB stack as well as light microscopy. Following alignment, numerous ARF1 stained tubules were inspected and all of them are correlated with corresponding tubular compartment in electron microscopy images. ARF1 compartments and clathrin coated compartments were segmented manually. Diameter of compartments and clathrin-coated membranes (Fig. 2D) was measured using the ruler tool in Dragonfly. The diameter of ARF1 compartments was measured in irregular distances within the ARF1 compartments shown in Fig. 2.

### Fixed-cell STED Z-stack

To preserve tubulo-vesicular structures for acquisition of 3D z-stack (Extended Data Fig. 5B), cells were fixed with 3% PFA and 0.2% glutaraldehyde, as described for FIB-SEM CLEM fixation. The z-stack was recorded with 200 nm increments.

### Trafficking assays

For the RUSH assay (Fig. 7A-F, H, Extended Data Fig. 6), plasmids encoding for the different RUSH cargoes as described in the figure legend were transiently transfected into HeLa cells and seeded on fibronectin coated microscopy dishes. 18 h after transfection, cells were stained with live-cell imaging dyes. The biotin stock solution (c=585 mM in DMSO) was diluted to a final concentration of 500 µM (in live-cell imaging solution) before addition to the cells. Live-cell confocal imaging was started 15 min, 20 min, 35 min or 70 min post biotin addition to investigate post-Golgi trafficking of the cargo.

For the Tfn assays (Fig. 8A, D-F, Extended Data Fig. 7), live-cell labeled HeLa cells were put on ice for 10-15 min before incubated with Tfn-AF488 (25 µM, ThermoFisher) in live-cell imaging solution at 37°C for 3 min. Cells were washed once with live-cell imaging solution before live-cell imaging with holo-Tfn (1 mM, ThermoFisher) was added to prevent re-endocytosis of labeled Tfn.

For the Tfn assays (Fig. 8C), live-cell labeled HeLa cells were put on ice for 10 min before incubated with Tfn-AF488 (25 µM, ThermoFisher) in live-cell imaging solution at 37°C for 2 min. For collecting the 2 min timepoint, cells were immediately washed 3x with PBS and then fixed with 3% PFA and 0.2% glutaraldehyde, as described for FIB-SEM CLEM fixation. For the later timepoints (5/7/10/15 minutes), cells were washed once with live-cell imaging solution before live-cell imaging with holo-Tfn (1 mM, ThermoFisher) was added to prevent re-endocytosis of labeled Tfn until the indicated timepoint. Cells were then washed 3x with PBS and fixed. Finally, the samples were mounted using pro-long Gold.

For the simultaneous RUSH and Tfn uptake assay (Fig. 8B), cells were prepared for the RUSH assay as described above. Live-cell imaging solution containing biotin was added to the cells (final concentration biotin= 500 µM) for 12 minutes at 37°C allowing it to pulse the secretory RUSH cargo out of the ER. Then, the medium was changed to live-cell imaging solution containing Tfn-AF488 (final concentration= 50 µM) for 30 s at 37°C allowing Tfn entry into the cell followed by live-cell imaging.

### Image Quantification

Prism was used to generate all graphs and to calculate SD and SEM. Data sets containing continuous data from different biological replicates were presented as superplots^65^.

To estimate the association of clathrin with ARF1 compartments and AP-2 (Fig. 1C), all clathrin punctae were manually counted in 10 ARF1^EN^-eGFP/AP2µ^EN^-SNAP/Halo-CLCa^EN^ HeLa cells from 3 independent experiments and categorized in 3 categories depending on their colocalization with either ARF1 or AP-2 or neither of them. A fourth category (ARF1+AP-2) shows the percentage of clathrin puncta which association could not be unambiguously determined. From each cell, an image at the Golgi plane and an image at the apical plasma membrane was acquired and used for quantification. For quantification of ARF1 compartments with different adaptor identity (Fig. 3D) ARF1^EN^-eGFP/AP1µA^EN^-SNAP/AP3µA^EN^-Halo HeLa cells were analyzed manually. Compartments were categorized based on which AP was found associated with the structure.

To estimate the amount of association of different APs with the Golgi, dual color confocal images of ARF1^EN^-Halo/AP1µA^EN^-SNAP, ARF1^EN^-Halo/AP3µA^EN^-SNAP and ARF1^EN^-Halo/AP4µ^EN^-eGFP HeLa cells were used for the analysis in Fiji. A mask of the Golgi was drawn using the ARF1-signal as reference. Using the “Find Maxima”-Tool of Fiji the number of AP puncta in the Golgi area and in the entire cell measured to estimate the percentage of Golgi associated AP.

To quantify the correlation of clathrin with AP-1 and AP-3 (Fig. 4E) the Manders correlation coefficient was used. Confocal images of ARF1^EN^-eGFP/AP1µA^EN^-SNAP/Halo-CLCa^EN^, ARF1^EN^-eGFP/AP3µA^EN^-SNAP/Halo-CLCa^EN^ and ARF1^EN^-eGFP/AP1µA^EN^-SNAP/AP3µA^EN^-Halo HeLa cells were background subtracted, gaussian blurred and the cell was selected as ROI. After automatic thresholding with Fiji the Manders correlation coefficient was determined.

To estimate the change in clathrin recruitment upon AP1µA KO (Fig. 4G), confocal images of ARF1^EN^-Halo/SNAP-CLCa^EN^ and ARF1^EN^-Halo/SNAP-CLCa^EN^ AP1µA KO HeLa cells were analyzed. Each cell was imaged at the plasma membrane plane and the Golgi plane. In each cell the fluorescence intensity of 10 clathrin puncta which were associated with ARF1 compartments were measured and averaged. For normalization, the average intensity of 10 clathrin plaques at the plasma membrane was used.

The overlap of ARF1 compartments and different endosomal markers (Fig. 5B) was quantified through a Manders correlation coefficient. For this, dual color overview images of HeLa cells expressing ARF1^EN^-SNAP/Halo-SNX1^EN^/Rab6/7/11^EN^ (labeled with BG-JF_552_ and CA-JFX_650_) and ARF1^EN^-Halo/SNAP-Rab5^OE^ (labeled with BG-JFX_650_ and CA-JF_552_) and as a positive control ARF1^EN^-Halo HeLa cells (labeled with CA-JFX_650_ and CA-JF_552_) were used. The confocal images were background subtracted, Gaussian blurred and the ROI was selected by hand excluding the nucleus and Golgi. The Manders values were calculated with a CellProfiler^66^ script summarized in Supplementary Table 5.

To estimate the overlap of compartments exiting the Golgi positive for ARF1 and Rab6 (Extended Data Fig. 3B), live-cell confocal time lapses of ARF1^EN^-SNAP/Halo-Rab6^EN^ HeLa cells labeled with BG-JF_552_ and CA-JFX_650_ were taken at 10 s/frame imaging speed. Carriers emerging from the Golgi were counted manually from 3 different cells.

The correlation of ARF1, Rab11 and AP-1 in Extended Data Fig. 4D was investigated by live-cell confocal imaging of ARF1^EN^-eGFP/Halo-Rab11^EN^/AP1µA^EN^-SNAP HeLa cells labeled with CA-JFX_650_ and BG-JF_552_ by taking square confocal images of the cytosol. The images were background subtracted and gaussian blurred and thresholded with Fiji before determining the Manders correlation coefficient with the JaCOP Fiji PlugIn^67^. To avoid higher values due to coincidental overlap, the obtained Manders values were subtracted by the Manders value measured once one channel is rotated by 90 degrees revealing the normalized Manders values shown in Extended Data Fig. 4D.

For visualization and quantification of maturation events, cells were imaged at a frame rate of 1.5 s/frame (Fig. 6). For 11 maturation events, the total fluorescence intensity of ARF1 and Rab11 was measured using Fiji. The images were background subtracted and gaussian blurred, and the compartment was selected as ROI and the total fluorescence signal for each channel was measured for the ROI. The ROI was adapted for each frame. The data sets were overlayed and averaged based on the timing of the drop in ARF1 fluorescence signal.

To quantify the percentage of secretory tubules exiting the Golgi positive for ARF1 (Fig. 7C), RUSH assays were performed in ARF1^EN^-Halo HeLa cells transiently overexpressing Streptavidin-KDEL/TfR-SBP-SNAP labeled with CA-JF_552_ and BG-JFX_650_. Live-cell confocal time lapses were taken 20 minutes post biotin addition at 6 s/frame imaging speed. Tubules emerging from the Golgi were counted manually from 3 different cells.

To understand the directionality of secretory cargo transfer between ARF1 compartments and RE (Fig. 7F), RUSH assays of Streptavidin-KDEL/TfR-SBP-SNAP transiently overexpressed in ARF1^EN^-Halo or Halo-Rab11^EN^ HeLa cells were performed. Images were captured 15 min post biotin at 1 min/frame. Confocal images were background subtracted, gaussian blurred and the ROI was selected by hand excluding the nucleus and Golgi. The threshold was obtained for every channel with Fiji and then Manders correlation coefficient was determined with the JaCOP plugin.

For determining the Golgi export delay upon AP1µA KO (Fig. 7H), RUSH assays were performed in ARF1^EN^-Halo (control) or ARF1^EN^-Halo AP1µA KO HeLa cells transiently overexpressing Streptavidin-KDEL/TfR-SBP-SNAP and ManII-GFP (Golgi mask) labeled with BG-JFX_650_. Live-cell confocal overview images were taken 15 min post biotin at 1 min/frame. The ROI was selected by hand excluding the nucleus and the Pearson correlation coefficient over time was determined using the JaCOP plugin of Fiji.

To place ARF1 compartments along the endocytic recycling route (Fig. 8C), Tfn assays were performed in ARF1^EN^-Halo, SNAP-Rab5^OE^ (early endosomes), Halo-Rab11^EN^ (recycling endosomes), and Halo-Rab6^EN^ (negative control) HeLa cells labeled with CA/BG-JFX_650_. After the indicated timepoint, cells were fixed as described for FIB-SEM CLEM fixation and a total of 10 cells (per timepoint of each endosomal marker) from 2 biological replicates were captured as confocal overview images. Images were background subtracted and Gaussian blurred in Fiji and the Manders values were calculated with a CellProfiler^66^ summarized in Supplementary Table 6.

To quantify the effect of an AP1µA KO on Tfn trafficking (Fig. 8E), Tfn assays were performed in ARF1^EN^-Halo and ARF1^EN^-Halo AP1µA KO HeLa cells labeled with CA-JFX650. Tfn recycling was recorded every 2 min for 30 min and the Manders correlation coefficient of ARF1 and Tfn was determined for every time point.

### Antibodies and Dyes

All live-cell imaging dyes used in the study are indicated in the figure legends and were provided from the Lavis lab^68,69^. All antibodies used in this study are provided in Supplementary Table 4.

## Acknowledgments

This project was supported by the Deutsche Forschungsgemeinschaft (German Research Foundation) grants SFB958 (Project A25 to F.B.), TRR186 (Projects A20 to F.B., A08 to V.H and Z02 to A.Z. and H.E.) and DFG grant HA 2686/24-1 (V.H.). C.R. is supported by a Human Frontier Science Program (HFSP) early career award to F.B.. We thank Luke Lavis (Janelia Research Campus) for providing all live-cell dyes we used in this study and Alexey Butkevich (Max Planck Institute for Biophysical Chemistry) for providing the SiX-lysosomal probe. We are grateful to David Gerishlick, Gaelle Boncompain and Franck Perez for sharing plasmids and expertise for RUSH experiments. We thank Stefan Donat and Jan Schmoranzer of the AMBIO imaging center (Charité Berlin) for the technical assistance for spinning disk experiments and our lab manager Giorgia Carai for general technical assistance.

## Author contribution

A.S., P.A. and F.B. conceived the project. A.S., P.A., V.N., A. H., A.K., G.B., S.H., C.R., C.S.,

A.G., J.M., H.E., D.P. V.H. and F.B. designed and performed experiments. A.S., P.A., V.N.,

A.H., A.K., S.H., A.G., A.Z. and C.R. performed image analysis. A.S., P.A., V.N., A.H., A.K., S.H., C.R., C.S., E.Ö., P.L., A.G., E.F., J.M. generated plasmids and knock in/out cell lines. A.S., P.A. and F.B. wrote the manuscript with input from all authors.

## Data availability

The data presented in this publication is available from the authors upon reasonable request.

## Authors note

The authors declare no competing interests exist.

