## Extended Data for "ARF1 compartments direct cargo flow via maturation into recycling endosomes"

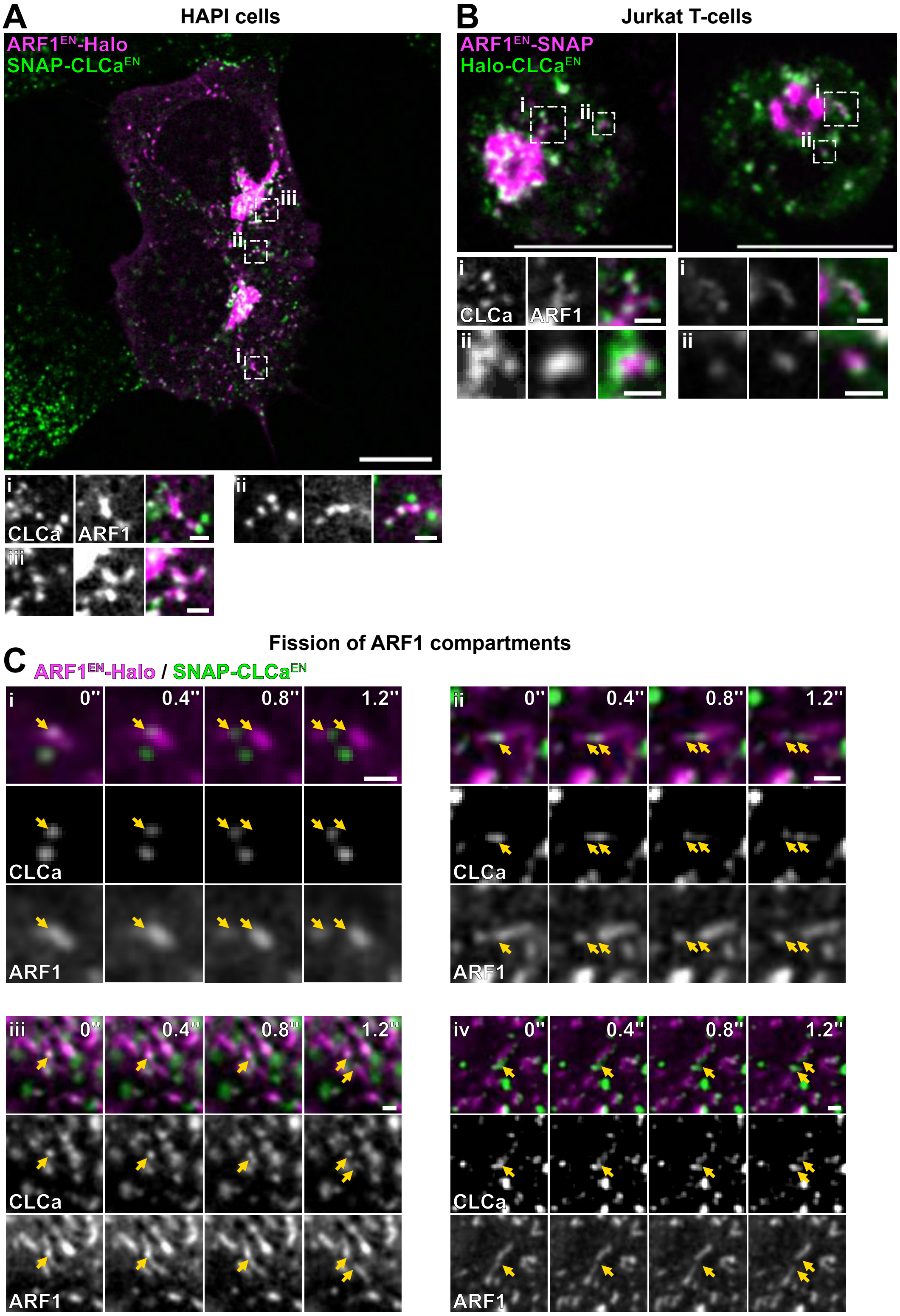

**Extended Data Figure 1: Clathrin is associated to ARF1 compartments in different cell types and fission occurs at sites of clathrin enrichment.** (**A-B**) Live-cell confocal imaging of ARF1$^{EN}$-Halo/SNAP-CLCa$^{EN}$ haploid HAP1 cells labeled with CA-JF$_{552}$ and BG-JFX$_{650}$ and ARF1$^{EN}$-SNAP/Halo-CLCa$^{EN}$ Jurkat T-cells labeled with CA-JF$_{571}$ and BG-JFX$_{650}$ highlight ARF1 compartments decorated by clathrin domains. (**C**) (i-iv) Multiple examples of time-lapse confocal spinning disk imaging of ARF1$^{EN}$-Halo/SNAP-CLCa$^{EN}$ HeLa cells labeled with CA-JF$_{552}$ and BG-JFX$_{650}$ showing clathrin at the site of fission on ARF1 compartments (yellow arrows highlight the localization of clathrin). Selected frames are shown, movie was taken with a frame rate of 5 frames/s. EN=endogenous, HaloTag substrate CA=chloroalkane, SNAP-tag substrate BG=benzylguanine. Scale bars: 10 µm (overviews) and 1 µm (crops).

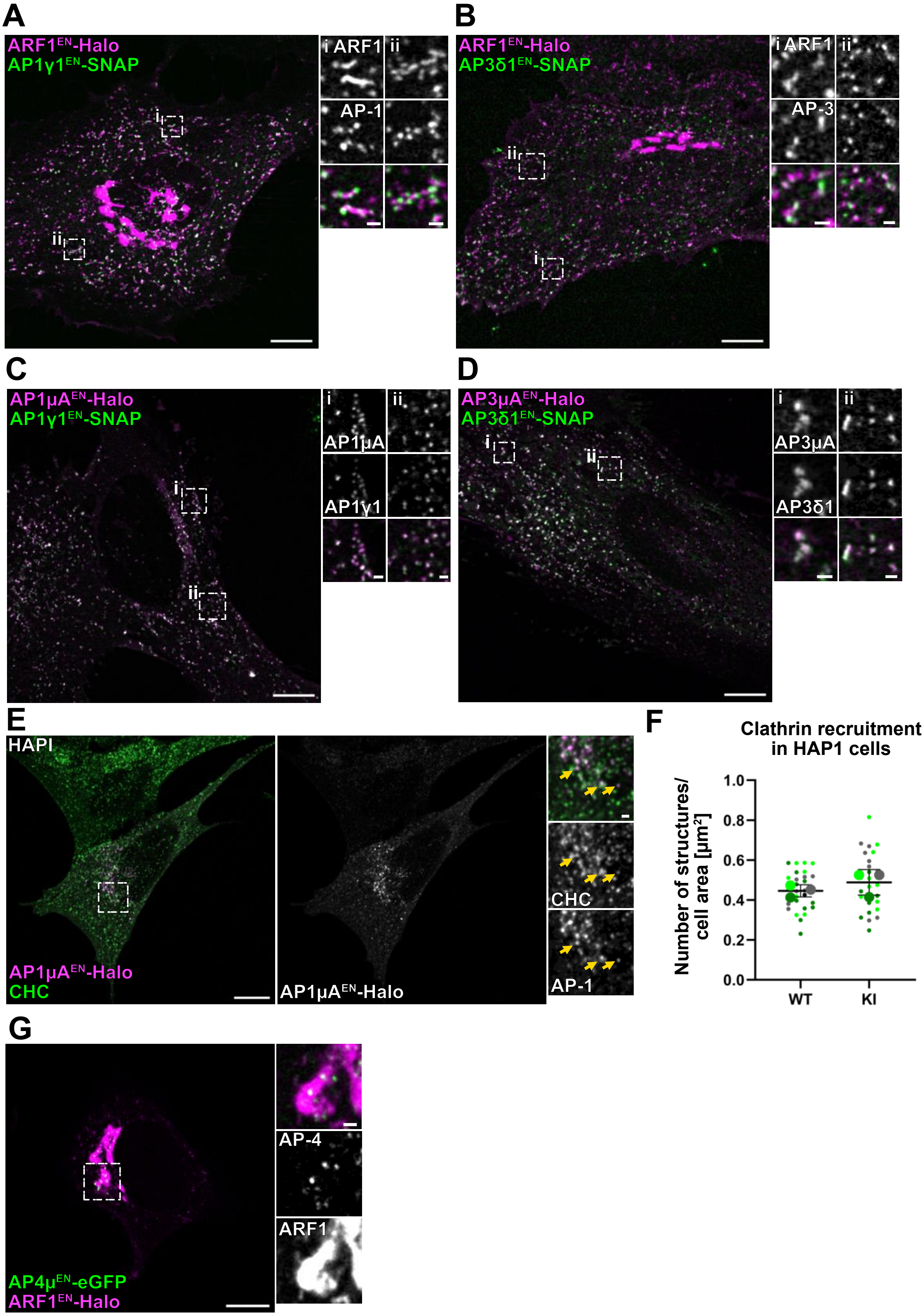
A
ARF1EN-Halo
AP1γ1EN-SNAP
i ARF1
ii
AP-1
B
ARF1EN-Halo
AP3δ1EN-SNAP
i ARF1
ii
AP-3
C
AP1μAEN-Halo
AP1γ1EN-SNAP
AP1μA
AP1γ1
D
AP3μAEN-Halo
AP3δ1EN-SNAP
AP3μA
AP3δ1
E
HAPI
AP1μAEN-Halo
CHC
AP1μAEN-Halo
CHC
AP-1
F
Clathrin recruitment in HAP1 cells
Number of structures/ cell area [μm²]
1.0
0.8
0.6
0.4
0.2
0.0
WT
KI
G
AP-4
ARF1
AP4μEN-eGFP
ARF1EN-Halo

**Extended Data Figure 2: Endogenous tagging of AP subunits does not affect AP function.** (**A-B**) Live-cell confocal imaging of ARF1$^{EN}$-Halo/AP1γ1$^{EN}$-SNAP and ARF1$^{EN}$-Halo/AP3δ1$^{EN}$-SNAP HeLa cells labeled with CA-JF$_{552}$ and BG-JFX$_{650}$ shows that C-terminal tagging of a large subunit of the adaptor complexes leads to comparable association with ARF1 compartments as observed for C-terminal tagging of the µ-subunit. (**C-D**) Live-cell confocal imaging of AP1µA$^{EN}$-Halo/AP1γ1$^{EN}$-SNAP and AP3µA$^{EN}$-Halo/AP3γ1$^{EN}$-SNAP HeLa cells labeled with CA-JF$_{552}$ and BG-JFX$_{650}$ shows that C-terminal tagging of two subunits of an APs has no effect on the localization of the complexes. (**E**) Confocal imaging of fixed AP1µA$^{EN}$-Halo HAP1 cells labeled with CA-JFX$_{650}$ before fixation and then immunostained with an anti-CHC (clathrin heavy chain) antibody. The recruitment of clathrin to AP-1 domains is unaffected by tagging of AP1µA (yellow arrows indicate domains where CHC and AP1µA$^{EN}$-Halo colocalize). (**F**) Quantification of clathrin puncta normalized to the cell area in wild type (WT) and AP1µA$^{EN}$-Halo knock-in (KI) HAP1 cells show no change in the number of cytosolic clathrin structures. In total 30 cells from 3 independent experiments for each condition were analyzed, replicates are shown in different colors and each dot represents a single cell. (**G**) Live-cell confocal imaging of ARF1$^{EN}$-Halo/AP4µ$^{EN}$-eGFP HeLa cells labeled with CA-JFX$_{650}$ showing AP-4 puncta at the TGN. EN=endogenous, HaloTag substrate CA=chloroalkane, SNAP-tag substrate BG=benzylguanine. Scale bars: 10 µm (overviews) and 1 µm (crops).

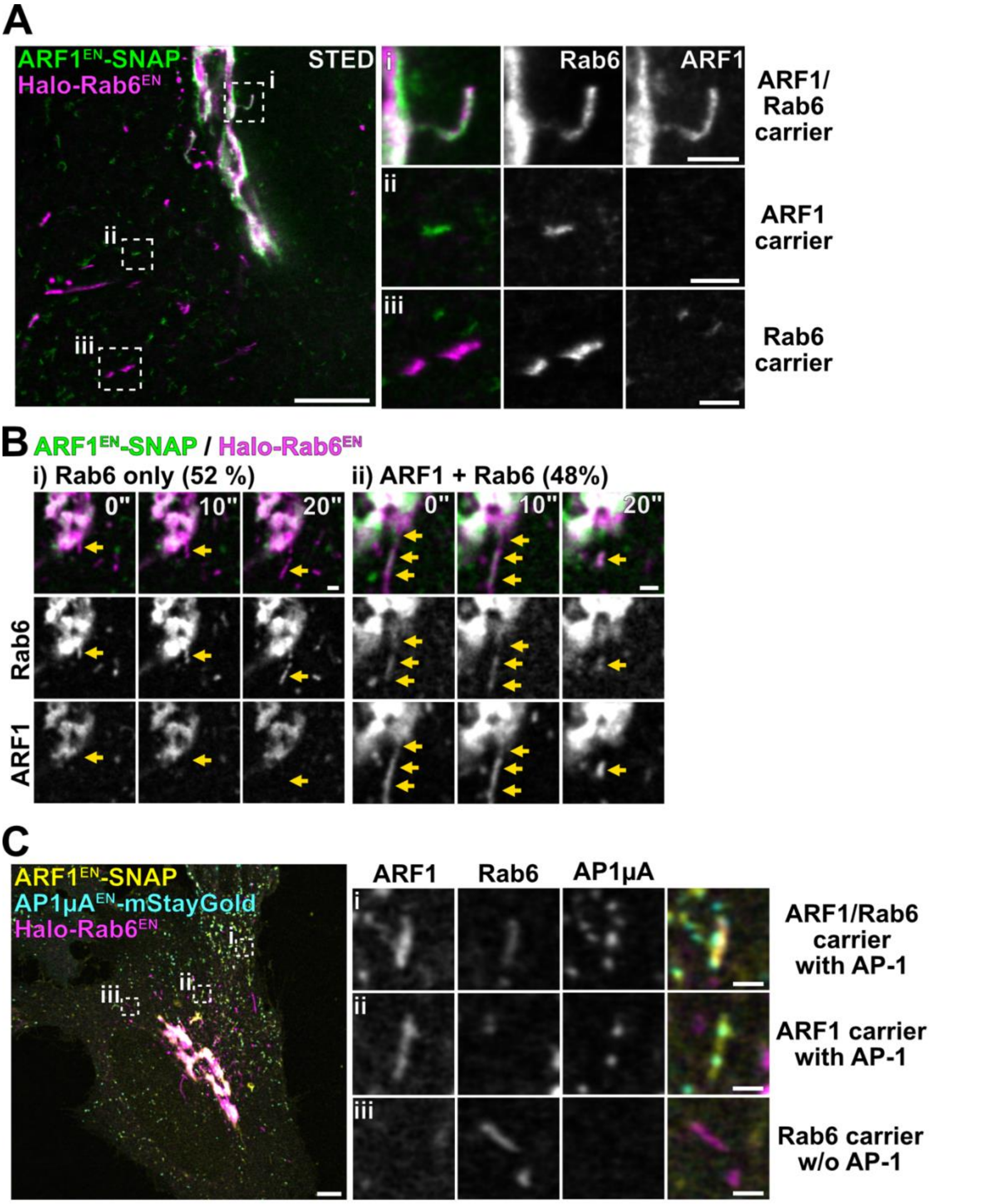


**Extended Data Figure 3: ARF1 defines a subpopulation of Golgi-derived Rab6 carriers.** (**A**) Live-cell STED imaging of ARF1$^{EN}$-SNAP/Halo-Rab6$^{EN}$ HeLa cells labeled with CA-JF$_{571}$ and BG-JFX$_{650}$ shows that (i) ARF1 and Rab6 on carriers that form at the TGN whereas (ii) peripheral ARF1 compartments are devoid of Rab6 and (iii) peripheral Rab6 carriers are devoid of ARF1. (**B**) Live-cell confocal imaging ARF1$^{EN}$-SNAP/Halo-Rab6$^{EN}$ HeLa cells labeled with BG-JF$_{552}$ and CA-JFX$_{650}$ shows that two distinct populations of Rab6 carriers emerging from the TGN: (i) half of the Rab6 carriers are devoid of ARF1 and (ii) half are marked by ARF1. (**C**) Live-cell confocal imaging of ARF1$^{EN}$-SNAP/Halo-Rab6$^{EN}$/AP1μA$^{EN}$-mStayGold HeLa cells labeled with BG-JF$_{552}$ and CA-JFX$_{650}$ shows that AP-1 only localizes to (i) ARF1/Rab6 double-positive carriers and (ii) ARF1 only but not to (iii) Rab6 only carriers. EN=endogenous, HaloTag substrate CA=chloroalkane, SNAP-tag substrate BG=benzylguanine. Scale bars: 5 µm (overview) and 1 µm (crops).

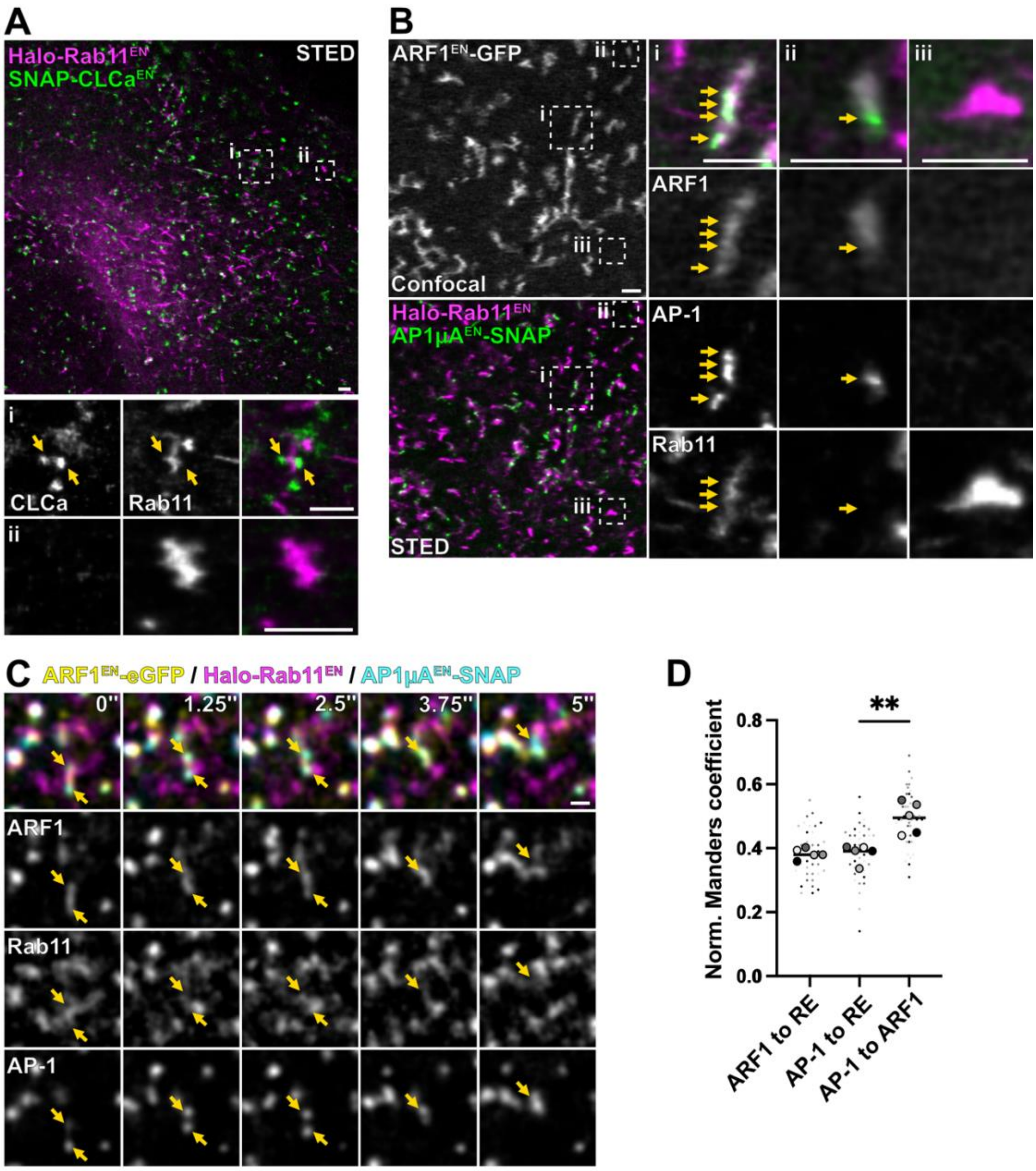


**Extended Data Figure 4: AP-1 and clathrin are recruited by ARF1 to RE membranes.** (**A**) Live-cell STED microscopy of $CLCa^{EN}$-SNAP/Halo-$Rab11^{EN}$ HeLa cells labeled with CA-$JFX_{650}$ and BG-$JF_{585}$ highlights (i) close proximity of clathrin coated structures to a RE and (ii) a RE devoid of clathrin. (**B**) Live-cell STED imaging of $ARF1^{EN}$-eGFP/Halo-$Rab11^{EN}$/$AP1\mu A^{EN}$-SNAP HeLa cells (two color STED imaging with $ARF1^{EN}$-eGFP imaged in confocal mode) labeled with BG-$JFX_{650}$ and CA-$JF_{571}$ shows (i) AP-1 on ARF1/Rab11 double-positive compartments, (ii) AP-1 on ARF1 compartments devoid of Rab11, (iii) Rab11-positive RE devoid of AP-1 and separated from ARF1 compartments. (**C**) Time-lapse confocal spinning disk imaging shows a moving ARF1 compartment that harbours AP-1 domains (yellow arrows) (**D**) Quantitative colocalization analysis between ARF1, AP-1 and RE. Normalized Manders coefficient shows comparable correlation of AP-1 to RE and ARF1 to RE, whereas correlation of AP-1 to ARF1 is significantly higher. Five replicates are shown in different colors each small dot representing a single cell of the replicate. P-value of the nested t-test is 0.0009. EN=endogenous, HaloTag substrate CA=chloroalkane, SNAP-tag substrate BG=benzylguanine. Scale bars: 1 µm.

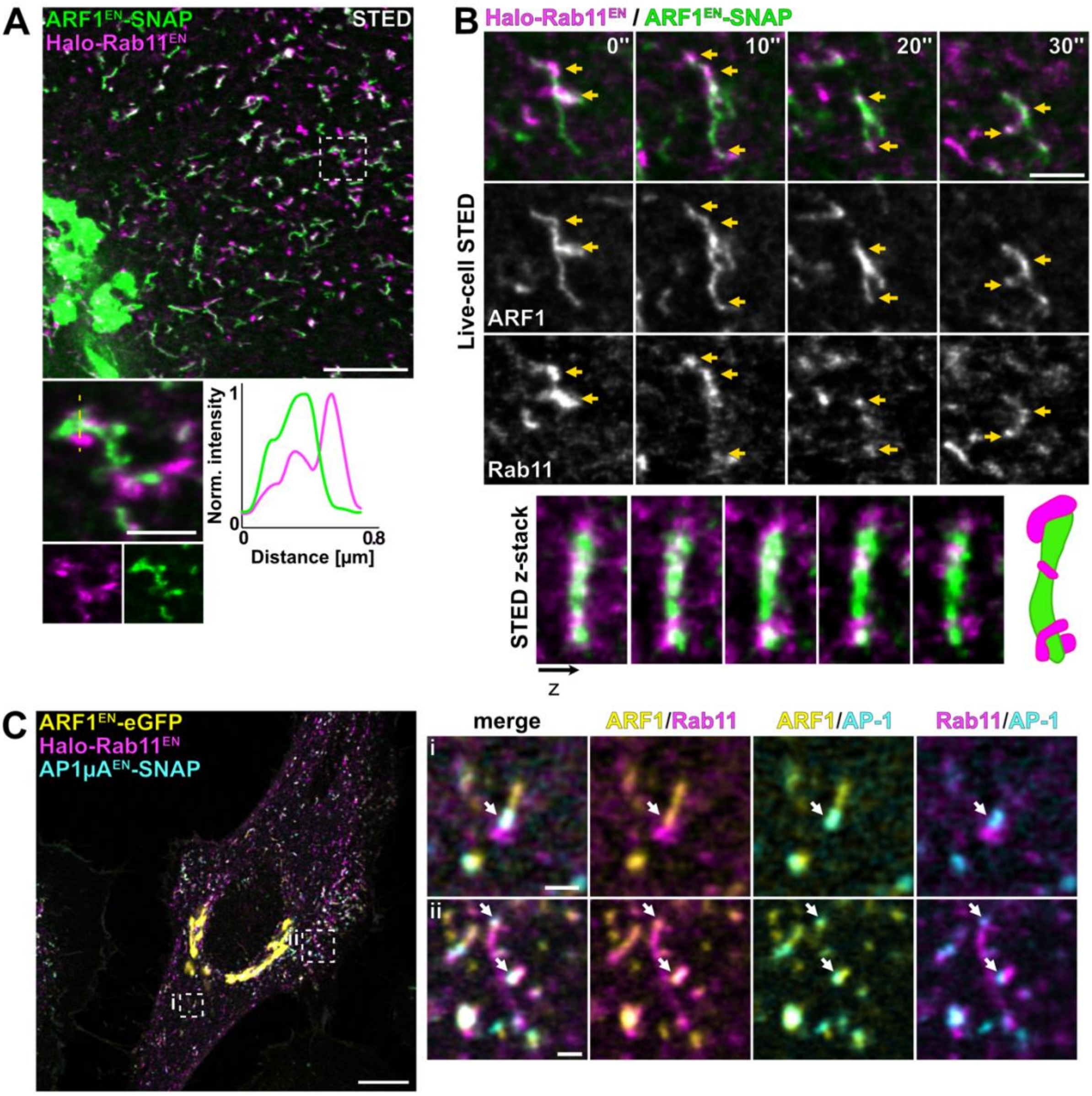


**Extended Data Figure 5: ARF1 compartments transiently interact with REs.** (**A**) Live-cell STED microscopy of ARF1$^{EN}$-SNAP/Halo-Rab11$^{EN}$ HeLa cells labeled with CA-JF$_{571}$ and BG-JFX$_{650}$ highlights ARF1 compartments and REs at nanoscale resolution. Line profile showing close association of ARF1 compartments and REs. (**B**) Live-cell STED imaging of ARF1$^{EN}$-SNAP/Halo-Rab11$^{EN}$ HeLa cells labeled with CA-JF$_{571}$ and BG-JFX$_{650}$ shows transient interaction of a peripheral ARF1 compartments and REs. Fixed-cell 3D-STED microscopy highlights close proximity of a single ARF1 compartment to multiple REs. (**C**) Live-cell confocal imaging of ARF1$^{EN}$-eGFP/Halo-Rab11$^{EN}$/AP1µA$^{EN}$-SNAP HeLa cells labeled with CA-JFX$_{650}$ and BG-JF$_{552}$ shows AP-1 at the interface of ARF1 compartment and RE located in the (i) periphery and (ii) perinuclear area of the cell. EN=endogenous, HaloTag substrate CA=chloroalkane, SNAP-tag substrate BG=benzylguanine. Scale bars: 10 µm (confocal overview), 5 µm (STED overview) and 1 µm (crops).

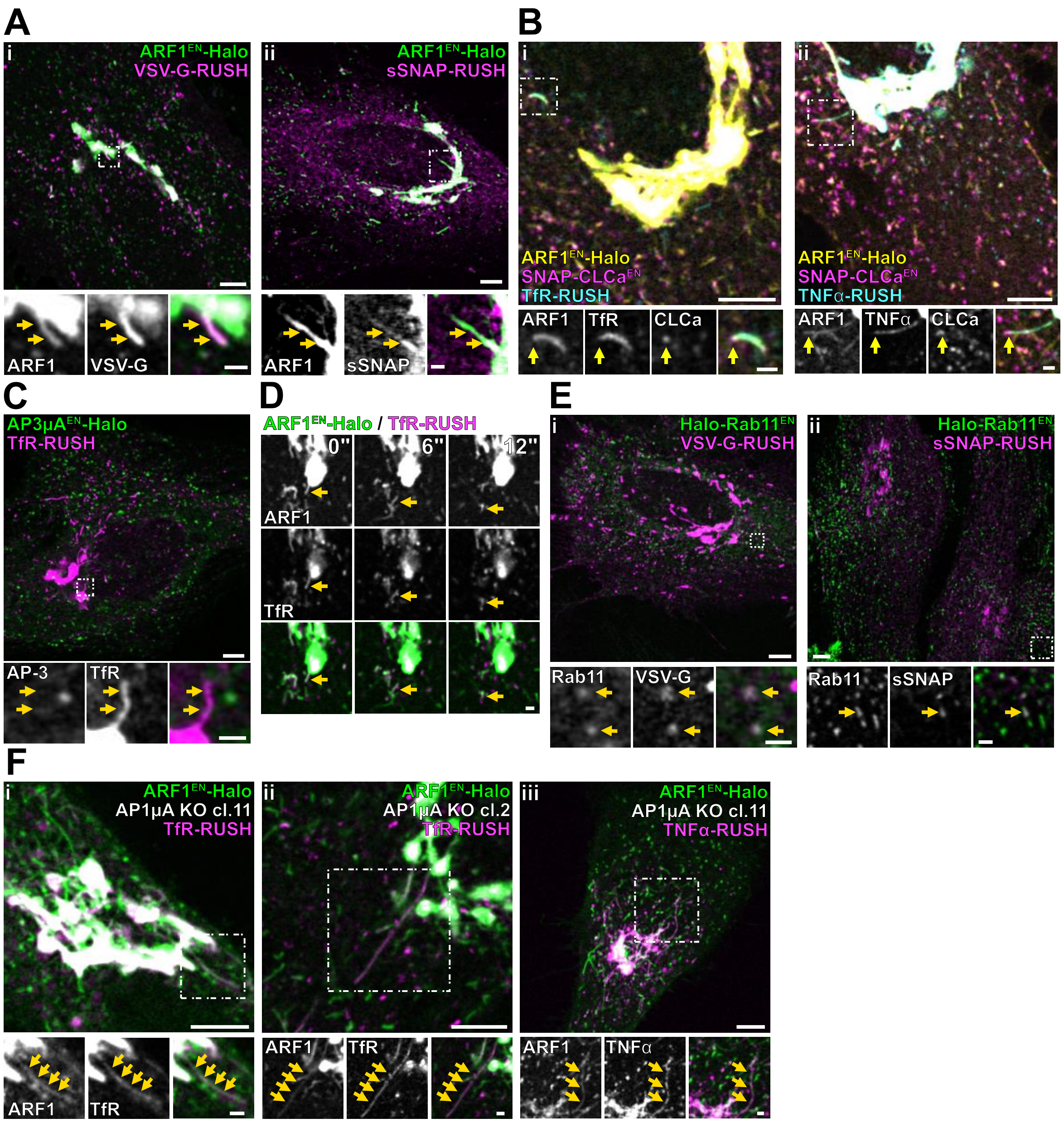


**Extended Data Figure 6: Perinuclear ARF1 compartments mediate secretory transport.** (**A**) Live-cell confocal imaging of ARF1$^{EN}$-Halo HeLa cells transiently expressing (i) Streptavidin-li/ssSBP-eGFP-VSV-G (labeled with CA-JFX$_{650}$) or (ii) Streptavidin-KDEL/ssSBP-SNAP (soluble SNAP (sSNAP), labeled with BG-JFX$_{650}$ and CA-JF$_{552}$) highlights the trans-membrane (VSV-G) and soluble RUSH cargo in ARF1 compartments. Image taken at 21 min (VSV-G) and 52 min (sSNAP) after addition of biotin. (**B**) Live-cell confocal imaging of ARF1$^{EN}$-Halo/SNAP-CLCa$^{EN}$ HeLa cells transiently expressing (i) Streptavidin-KDEL/TfR-SBP-eGFP or (ii) Streptavidin-KDEL/TNFα-SBP-eGFP labeled with BG-JFX$_{650}$ and CA-JF$_{552}$ highlights trans-membrane RUSH cargo (TfR, TNFα) in ARF1 compartments decorated by clathrin (yellow arrows highlight clathrin on ARF1 compartments). Images were taken (i) 20 min and (ii) 16 min after addition of biotin. (**C**) Live-cell confocal imaging of AP3μA$^{EN}$-Halo HeLa cells transiently expressing Streptavidin-KDEL/TfR-SBP-eGFP labeled with CA-JFX$_{650}$ show that RUSH carriers are devoid of AP-3. Image taken 23 min after addition of biotin. (**D**) Live-cell confocal time-lapses of ARF1$^{EN}$-Halo HeLa cells transiently expressing Streptavidin-KDEL/TfR-SBP-SNAP labeled with BG-JFX$_{650}$ and CA-JF$_{552}$ show ARF1 compartments decorated by TfR-RUSH detaching from the Golgi and moving away (highlighted by a yellow arrow). The first frame was taken at 23 min after addition of biotin. (**E**) Live-cell confocal imaging of Halo-Rab11$^{EN}$ HeLa cells transiently expressing (i) Streptavidin-li/ssSBP-eGFP-VSV-G (labeled with CA-JFX$_{650}$) or (ii) Streptavidin-KDEL/ssSBP-SNAP (labeled with BG-JFX$_{650}$ and CA-JF$_{552}$) highlights the trans-membrane (VSV-G) and soluble (sSNAP) RUSH cargo localization to recycling endosomes. Image taken at 25 min (VSV-G) and 70 min (sSNAP) after addition of biotin. **(F)** Live-cell confocal imaging of ARF1$^{EN}$-Halo/AP1μA KO HeLa cells transiently expressing (i) Streptavidin-KDEL/TfR-SBP-SNAP

(ii) Streptavidin-KDEL/TfR-SBP-eGFP or (iii) Streptavidin-KDEL/TNF$\alpha$-SBP-SNAP labeled with (i, iii) BG-JFX$_{650}$ and CA-JF$_{552}$ or (ii) CA-JFX$_{650}$ show trans-membrane RUSH cargo in aberrant elongated ARF1 compartments. (i-iii) Different CRISPR-Cas9 KO clones display the same phenotype. Images taken (i) 32 min or (ii) 25 min or (iii) 30 min after addition of biotin. EN=endogenous, HaloTag substrate CA=chloroalkane, SNAP-tag substrate BG=benzylguanine, ss=signal sequence.
Scale bars: 5 µm (confocal overviews), 1 µm (crops).

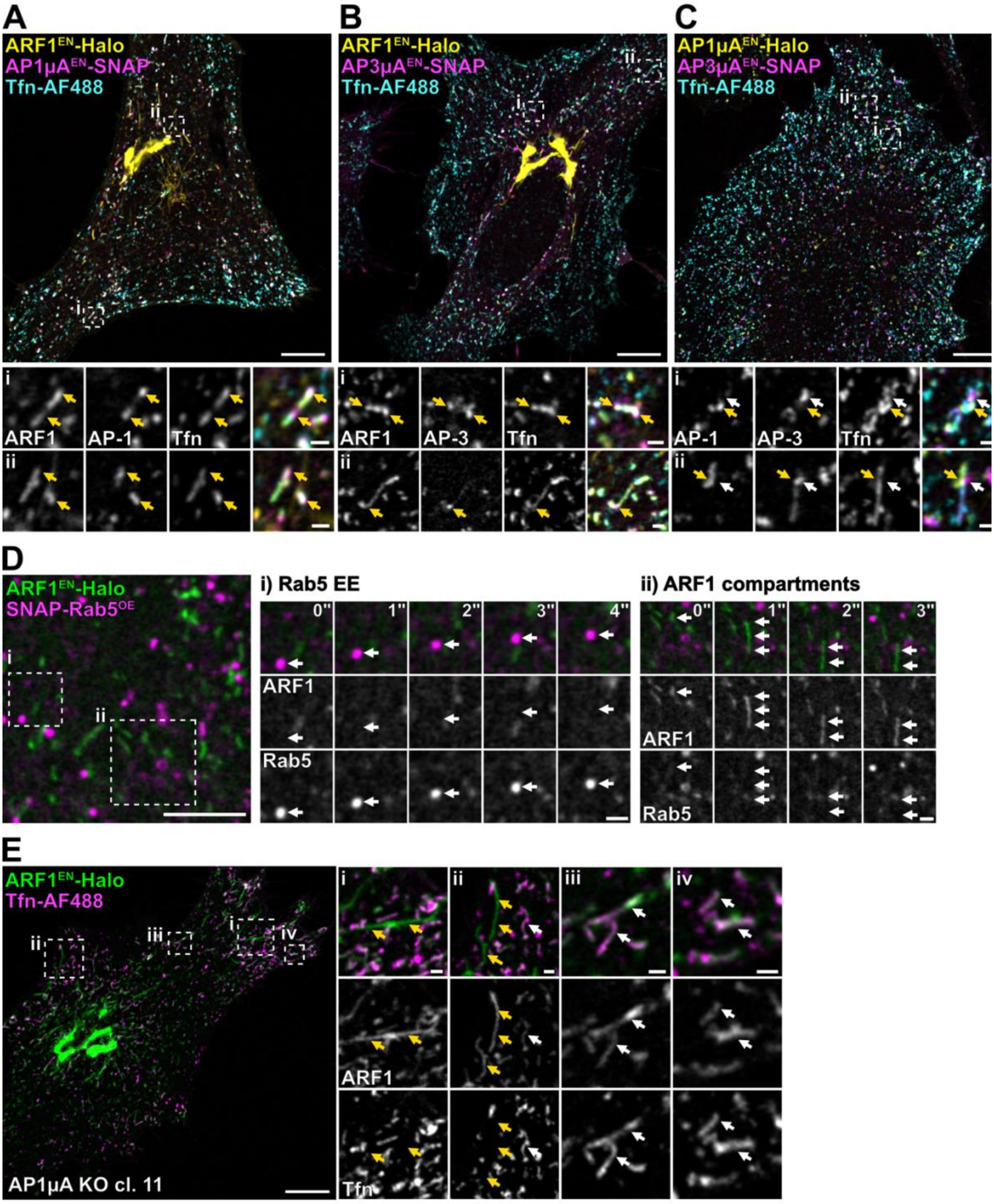


**Extended Data Figure 7: Peripheral ARF1 compartments mediate endocytic recycling of Tfn and are not derived from early endosomes.** Live-cell confocal imaging of fluorescently labeled Tfn (Tfn-AlexaFluor488) in different KI and KO HeLa cells. (**A-B**) ARF1$^{EN}$-Halo/AP1μA$^{EN}$-SNAP and ARF1$^{EN}$-Halo/AP3μA$^{EN}$-SNAP HeLa cells labeled with CA-$JF_{552}$ and BG-$JFX_{650}$. Fluorescent transferrin localizes to ARF1 compartments harbouring AP-1 and AP-3 domains (yellow arrows). (**C**) AP1μA$^{EN}$-Halo/AP3μA$^{EN}$-SNAP HeLa cells labeled with CA-$JF_{552}$ and BG-$JFX_{650}$. Fluorescent transferrin localizes to ARF1 compartments harbouring AP-1 (yellow arrows) and AP-3 (white arrows) domains. (**D**) Live-cell confocal imaging of gene edited ARF1$^{EN}$-Halo HeLa cells transiently overexpressing SNAP-Rab5$^{OE}$ labeled with CA-$JF_{503}$ and BG-$JFX_{650}$. ARF1 compartments are not derived from early endosomes (EE) and (i) Rab5 positive EE and (ii) ARF1 compartments move independently. (**E**) ARF1$^{EN}$-Halo/AP1μA KO HeLa cells labeled with CA-$JFX_{650}$. Tfn fills ARF1 compartments which morphology is unaffected upon loss of AP1μA. (i-ii) yellow arrows highlight aberrant elongated ARF1 compartments devoid of Tfn. (ii-iv) white arrows highlight shorter ARF1 compartments filled with Tfn. EN=endogenous, HaloTag substrate CA=chloroalkane, SNAP-tag substrate BG=benzylguanine. Scale bars: 10 µm (A-C, E overviews), 5 µm (D overviews), 1 µm (crops).
