## Supplementary Methods for "ARF1 compartments direct cargo flow via maturation into recycling endosomes"

For all plasmids the used HaloTag originates from pH6HTC His6HaloTag® T7 (Promega, JN874647) and the SNAP-tag from pSNAPf (New England Biolabs, N9183S). All plasmids were verified through sequencing. Sequences of all primers are provided in Supplementary Table 1.

**Generation of overexpression (OE) plasmids**

For generating SNAP-Rab5A, the Rab5 fragment was amplified from eGFP-Rab5A (addgene plasmid #49888) and cloned into a pSNAPf backbone using BamHI and NotI.

For the generation of SNAP-tagged RUSH cargoes, Streptavidin-KDEL/TNFα-SBP-eGFP and Streptavidin-KDEL/TfR-SBP-eGFP were used^1^. eGFP was exchanged to SNAP (Streptavidin-KDEL/TfR and TNFα-SBP-SNAP) by amplifying SNAP as a Sbfl/Xbal fragment from pSNAPf and inserting it into the Sbfl/Xbal linearized eGFP plasmids.

For generating the soluble SNAP (sSNAP) cargo (Streptavidin-KDEL/ss(signal sequence)-SBP-SNAP), a ssSBP-SNAP fragment was synthesized as a gBlock and inserted into the Xbal/EcoRI linearised Streptavidin-KDEL/SBP-SNAP-colX vector^1^. For creating Streptavidin-KDEL/ssSBP-SNAP-LAMP1Δ (w/o QYTI), the vector Str-KDEL-SBP-SNAP-CollagenX was PacI/Xbal digested and SNAP-LAMP1Δ was inserted as gBlock through Gibson assembly. ManII-eGFP^2^ and Streptavidin-li/ssSBP-eGFP-VSV-G^1^ were described previously.

**Generation of CRISPR knock-in (KI) cell lines**

**Cloning of guide and homology repair (HR) plasmids**

All guide RNAs were designed using the online tool Benchling (https://www.benchling.com) and cloned into either the SpCas9 pX459 plasmid (addgene plasmid #62988)^3^ or the SpCas9 pX330 plasmid (addgene plasmid #42230)^4^ by annealing oligos and ligation into the vector which was linearized with BbsI. A detailed list of guide RNA sequences is provided in Supplementary Table 2.

**Cloning of HR plasmids**

To avoid re-cutting from the Cas9, the protospacer-adjacent motif (PAM) site was mutated in the HR plasmids (if part of the coding region a silent mutation was introduced). The G418 resistance (G418^R^ containing SV40 promoter, ORF, polyadenylation signal) cassette originates from pEGFP-N1 (Clontech), the Puromycin resistance cassette (Puro^R^ containing SV40 promoter, ORF, polyadenylation signal) from pPUR (Clontech), the Hygromycin resistance cassette (Hygromycin^R^ containing SV40 promoter, ORF, polyadenylation signal) from pcDNA5/FRT/TO V5 (addgene plasmid #19445^5^) and the SV40 polyadenylation signals (PolyA) sequence originates from pEGFP-N1.

*HR plasmids AP1µA:*

The HR plasmid was designed with ~1Kb homology arms and was synthesized by Twist Bioscience. A glycin-serin linker and a BamHI and EcoRI site were added between the two homology arms for insertion of tags and resistance cassette. The coding sequences of HaloTag, SNAP-tag and mStayGold including a small epitope-tag were integrated between the homology arms, followed by a polyA sequence and a G418^R^ (in combination with HaloTag),a Puro^R^ (in combination with SNAP-tag) or a Hygromycin^R^ (in combination with mStayGold). The coding sequences of HaloTag, SNAP-tag were obtained via PCR using sense primers with a BamHI restriction site and antisense primers with a NheI restriction site. The SNAP-tag antisense primer included the sequence of the V5 epitope tag. The SV40 polyA sequence was amplified from pEGFP-N1 using a PolyA NheI sense and a PolyA NotI for pEGFP-C1 using a G418 NotI sense and G418 EcoRI antisense primer. The Puro^R^ cassette was amplified using a Puro NotI sense and Puro EcoRI antisense primer.. The various fragments were cloned into the HR vector linearized with BamHI and EcoRI. To generate the AP1µA^EN^-mStayGold-PolyA-Hygromycin^R^ HR plasmid, the AP1µA^EN^-Halo-PolyA-G418^R^ HR plasmid was digested with BamHI and NheI and the coding sequence of mStayGold was inserted as gBlock to replace the Halo coding sequence. In a second step the Hygromycin^R^ cassette was amplified using a Hygromycin ClaI sense and Hygromyicin EcoRI antisense primer and inserted into the AP1µA-StayGold HR vector linearized with ClaI and EcoRI.

*HR plasmid AP2µ:*

The HR plasmid was designed according to the design of the HR plasmid for AP1µA and was synthesized by Twist Bioscience. To generate the AP2μ^EN^-SNAP-V5-PolyA-Puro^R^ HR plasmid the entire insert (SNAP-V5-PolyA-Puro^R^) was excised from the AP1μA^EN^-SNAP-V5-PolyA-Puro HR plasmid using the BamHI and the EcoRI site and inserted into the ordered AP2µ plasmid, linearized with BamHI and EcoRI.

*HR plasmids AP3µA:*

The HR plasmid was designed according to the design of the HR plasmid for AP1µA and was synthesized by Twist Bioscience. To generate the AP3μA^EN^-SNAP-V5-PolyA-Puro HR plasmid the entire insert (SNAP-V5-PolyA-Puro^R^) was excised from the AP1μA^EN^-SNAP-V5-PolyA-Puro^R^ HR plasmid using the BamHI and the EcoRI site and inserted into the ordered AP3µA plasmid, linearized with BamHI and EcoRI. To generate the AP3μA^EN^-Halo-ALFA-PolyA-G418^R^ HR plasmid the PolyA-G418 fragment was excised from AP1μA^EN^-Halo-PolyA-G418^R^ HR using NheI and EcoRI. The coding sequences of HaloTag was obtained via PCR using a sense primer with a BamHI restriction site and an antisense primer with a NheI restriction site which included the sequence of the ALFA epitope tag. Both fragments were inserted into the ordered AP3µA HR plasmid, linearized with BamHI and EcoRI.

*HR plasmids AP4µ:*

The HR plasmid was designed according to the design of the HR plasmid for AP1µA and was synthesized by Twist Bioscience. To generate the AP4μ^EN^-eGFP-PolyA-G418^R^ HR plasmid the PolyA-G418^R^ fragment was excised from AP1μA^EN^-Halo-PolyA-G418^R^ HR using NheI and EcoRI. The coding sequence of eGFP was obtained via PCR from pEGFP-C1 using a sense primer with a BamHI restriction site and an antisense primer with a NheI restriction site.

*HR plasmids AP1γ1 and AP3δ1:*

The HR plasmids were designed according to the design of the HR plasmid for AP1µA and was synthesized by Twist Bioscience with homology arms shortened to 500 bp. To generate the AP1γ1^EN^-SNAP-V5-PolyA-Puro^R^ and AP3δ1^EN^-SNAP-V5-PolyA-Puro^R^ HR plasmid the entire insert (SNAP-V5-PolyA-Puro^R^) was excised from the AP1μA^EN^-SNAP-V5-PolyA-Puro^R^ HR plasmid using the BamHI and the EcoRI site and inserted into the ordered HR plasmids, linearized with BamHI and EcoRI.

*HR plasmids CLCa:*

The HR plasmid designs of LoxP-Puro^R^-LoxP-SNAP-CLCa^EN^ and LoxP-G418^R^-LoxP-Halo-CLCa^EN^ were previously described^6^.

*HR plasmids ARF1:*

The design of ARF1^EN^-Halo/SNAP-PolyA-G418^R^ HR plasmid was previously described^7^. To generate the ARF1^EN^-eGFP HR plasmid an alternative version of the HR plasmid was synthesized by Twist Bioscience including ~1Kb homology arms with a glycine-serin linker and a NheI and BamHI site between the two homology arms. The coding sequences of eGFP was obtained via PCR from pEGFP-C1 using sense primers with a NheI restriction site and antisense primers with a BamHI restriction site. The fragment was cloned into the HR plasmid linearized with NheI and BamHI.

To generate the ARF1^EN^-SNAP-V5-PolyA-Puro^R^ HR plasmid, first a SNAP-V5-tag fragment was obtained via PCR using a sense primer and antisense with a NheI restriction site and inserted in the ARF1 HR plasmid linearized with NheI. In a second step, the PolyA-Puro^R^ fragment was obtained via PCR using a sense with a NheI site and antisense primer with a BamHI restriction site and inserted in the ARF1^EN^-SNAP-V5 HR plasmid linearized with NheI and BamHI. To generate the ARF1^EN^-mStayGold HR plasmid, an alternative version of the HR plasmid was synthesized by Twist Bioscience including 500 bases homology arms with a glycine-serin linker and a BamHI and EcoRI site between the two homology arms. A mStayGold-PolyA-Hygromycin^R^ fragment was excised from the AP1μA^EN^-mStayGold-PolyA-Hygromycin^R^ HR plasmid using BamHI and the EcoRI and inserted into the new ARF1 HR plasmid, linearized with BamHI and EcoRI.

*HR plasmids Rab11A and Rab7A:*

The HR plasmid for endogenously tagging Rab11A N-terminally, was designed with ~1000 bp homology arms based on the region around the start codon. The HR plasmid was constructed starting from pHaloSec61ß plasmid, which is based on pEGFP-C1 and contains a Halo tag. The homology arms were amplified from genomic wild-type HeLa DNA. First, the left homology arm (LHA) was cloned using Asel and NheI and then the right homology arm (RHA) using MluI and BglII. A GS linker is included before the RHA. Lastly, a G418^R^ flanked by loxP sites was cloned into the full HR plasmid as a NheI fragment. The HR plasmid for endogenously tagging Rab7A N-terminally was generated like Rab11A with the exception that the homology arms were ordered as G-blocks and inserted through Gibson assembly at the AseI/NheI (LHA) and MluI/BglII (RHA) site.

*HR plasmids Rab6 and SNX1:*

The HR plasmids for endogenously tagging Rab6 and SNX1 N-terminally, were designed with ~700 bp homology arms based on the region around the start codon and synthesized by GeneScript (SNX1) or Thermofisher Scientific (Rab6) containing a GS linker located before the RHA. The designed HR plasmids contain the restriction sites NheI/SpeI (SNX1) or NheI/BamHI (Rab6) for insertion of tags and additional DNA between the two homology arms. Generally, for addition of a N-terminal tag, a plasmid was constructed in the lab containing a G418^R^ flanked by loxP sites, 3xALFA tag (ordered as a G-Block from IDT and inserted through Gibson assembly) and a Halo tag containing NheI/SpeI. This loxP-G418^R^-loxP-3xALFA-Halo fragment was excised and cloned into the synthesized SNX1 HR linear plasmid at the NheI/SpeI site. For inserting the N-terminal tag into the synthesized Rab6 HR plasmid the loxP-G418^R^-loxP-3xALFA-Halo was amplified as a NheI/BamHI fragment and inserted into the NheI/BamHI linearized Rab6 HR plasmid.

**Generation of CRISPR KI cell lines**

For generation of HeLa KI cell lines, cells at 70-80% confluency were transiently transfected with both guide and HR plasmids using FuGENE HD (Promega) according to the supplier’s protocol.

For generation of HAP1 KI cell lines, 1 million HAP1 cells were washed twice with Opti-MEM, resuspended in 90 µl Opti-MEM and mixed with 5 µg of guide and HR plasmid DNA in an electroporation cuvette with a 2-mm gap. The electroporation reaction consists of two poring pulse (125 V, 3 ms length, 50 ms interval, with decay rate of 10% and + polarity) and five consecutive transfer pulses (20 V, 50 ms length, 50 ms interval, with a decay rate of 40% and ± polarity).

In both cases, G418, puromycin or hygromycin were added to the cells 3 days after transfection at a concentration of 1.5 mg/mL (G418) or 2 µg/mL (puromycin) or 4 mg/ml (hygromycin) and media was exchanged every 2-3 days until selection was complete (G418: 7-10 days, puromycin: 2-5 days, hygromycin 6-8 days). After the selection of N-terminal KIs, cells were again transfected with a Cre-recombinase (addgene plasmid #11923)^8^ using FuGENE HD to remove the loxP-flanked resistance cassette.

For generation of KI Jurkat cell lines, 5 million Jurkat T cells were washed twice with Opti-MEM (Gibco), resuspended in 90 µl Opti-MEM and mixed with 2.5 µg of the guide and HR plasmids DNA in an electroporation cuvette with a 2-mm gap. The electroporation reaction consists of two poring pulse (150 V, 5 ms length, 50 ms interval, with decay rate of 10% and + polarity) and five consecutive transfer pulses (20 V, 50 ms length, 50 ms interval, with a decay rate of 40% and ± polarity). G418 was added to the cells 4 days after transfection at a concentration of 3 mg/mL and media was exchanged every 2-3 days for 7 days. Cells were again electroporated using the same protocol with a Cre-recombinase (addgene plasmid #11923)^8^ to remove the loxP-flanked resistance cassette. After 3 days, G418 was added to the cells at a concentration of 3 mg/ml media and was exchanged every 2-3 days for 5 days.

All KI cell lines were validated via western blotting. An overview of plasmids, base cell lines and selection method used for creation of new KI cell lines is given in Supplementary Table 3.

**Generation of AP1µA knock-out (KO) cell lines**

To achieve an AP1µA KO, the AP1M1 gene was targeted with to guide RNAs binding to sequences in exon 2 and exon 5. Guides were designed with the online tool Benchling (https://www.benchling.com) and cloned into the SpCas9 pX459 plasmid (addgene plasmid #62988)^3^ by annealing oligos and ligation into the vector which was linearized with BbsI. The KO guide RNA sequences are provided in Supplementary Table 2.

For generation of AP1µA KO cell lines, cells at 70-80% confluency were transiently transfected with both guide plasmids using FuGENE HD (Promega) according to the supplier’s protocol. One day after transfection, transfected cells were selected with 2 µg/mL of puromycin. Single cell clones were obtained via serial dilution. KO of AP1µA in single cell clones was confirmed via western blot.

An overview of plasmids, base cell lines used for creation of AP1µA KO cell lines is given in Supplementary Table 3.

**SDS page and western blot**

Cells lysates were loaded on 4-12% SDS-polyacrylamide gels (Life Technologies) and after electrophoresis, proteins were transferred to a nitrocellulose membrane (Amersham) via wet blotting. Membranes were blocked with 5% (wt/vol) milk powder and 1% BSA in PBST and incubated with primary antibodies overnight. For detection a secondary horseradish peroxidase-coupled antibody to the primary antibody was used. To develop the membrane the ECL western blot substrate was added for 2 min and then the membrane was imaged. Used antibodies are listed in Supplementary Table 4.

**Immunostaining of AP1µA-Halo HAP1 cells**

AP1µA-Halo HAP1 cells were seeded on fibronectin-coated coverslips and on the next day live-cell labelled with CA-JFX_650_. After washing the cells were rinsed twice with PBS and fixed with 4% PFA for 10 min. Samples were rinsed three time with PBS and incubated in permeabilization buffer (0.3% NP40, 0.05% Triton-X 100 and 0.1% BSA (IgG free) in PBS) for 3 min. Following this, samples were blocked for 1 h in blocking buffer (0.05% NP40, 0.05% Triton-X 100 and 5% goat serum (Jackson ImmunoResearch) in PBS), incubated with primary anti-CHC antibody (1:1000 in blocking buffer) overnight and washed three times with washing buffer (0.05% NP40, 0.05% Triton-X 100 and 0.2% BSA in PBS). Finally, samples were incubated for 1 h with Alexa594-labelled secondary antibody (1:5000 in blocking buffer), washed three times 5 min with washing buffer and mounted with pro-long Gold (Life Technologies).

**Supplementary Table 1: Primers**

| Primer Name | Primer Sequence (5’-3’) |
| --- | --- |
| Rab5A-SNAP OE plasmid: | |
| Rab5A BamHI sense | CGAAGGGATCCGG ATCAGGTTCTATG GCTAGTCGAGGCG CAAC |
| Rab5A NotI antisense | CCGTAGCGGCCG CTTAGTTACTACA ACACTGAT |
| Streptavidin-KDEL/TfR or TNFα-SBP-SNAP OE plasmid: | |
| SNAP SbfI sense | ATAAcctgcaggtGACAAAGACTGCGAAATGAAG |
| SNAP XbaI antisense | TTAAtctagaTTAACCCAGCCCAGGCTTGCCCAG |
| Streptavidin-KDEL/SBP-sSNAP | |
| SBP EcoRI sense | TCACGAATTCCGACGAGAAGACCACTGGTTGGCGAGGTG |
| SNAP XbaI antisense | TGACTCTAGATTAACCCAGCCCAGGCTTGCCCAGTCTGTGGCC |
| Rab11A homology arms: | |
| Rab11A RHA BglI sense | TGCTAAGATCTGGATCAGGGTCAGGCTCTGGTTCCG GCTCAATGGGCACCCGCGACGACGAATATGACTACC TCTTTAAAGGTG |
| Rab11A RHA MluI antisense | AGTCAACGCGTGTATTTGTGCAGCGCACAACCTTTCC TGC |
| Rab11A LHA AseI sense | TGCAGATTAATGTATCCAAACTTCCGCTAAAATGGAAA TTACC |
| Rab11A LHA NheI antisense | TCGTAGCTAGCTGCGCTGCCGAGGAGCGAAAGGGC GGGAGCAGC |
| loxP-G418^R^-loxP-3xALFA-Halo as a NheI/SpeI fragment for N-terminal KI: | |
| loxP-G418^R^-loxP NheI sense | ATTACGCTAGCATAACTTCGTATAGCATACATTATACGAAGTTATCCTGAGGCGGAAAGAACCAGCTGTGGAATGTGTGTCAGTTAG |
| loxP-G418^R^-loxP XhoI antisense | ATTACCTCGAGATAACTTCGTATAATGTATGCTATACGAAGTTA TTTTATTCTGTCTTTTTATTGCCGTCATAGCGCGGGTT |
| 3xALFA Halo XhoI sense | TATTGCTCGAGGTATCATCTAACGGATCGGTATGAG |
| Halo SpeI antisense | TTAGCACTAGTGCCGGAAATTTCGAGCGTCGACAGC |
| loxP-G418^R^-loxP-3xALFA-Halo as a NheI/BamHI fragment for N-terminal KI | |
| loxP-G418^R^-loxP NheI sense | ATTACGCTAGCATAACTTCGTATAGCATACATTATACGAAGTTATCCTGAGGCGGAAAGAACCAGCTGTGGAATGTGTGTCAGTTAG |
| Halo BamHI antisense | cgaagGGATCCGCCGGAAATCTCGAGCGTCG |
| loxP-G418-loxP fragment for N-terminal KI: | |
| loxP-G418-loxP NheI sense | ATTACGCTAGCATAACTTCGTATAGCATACATTATACGAAGTTATCCTGAGGCGGAAAGAACCAGCTGTGGAATGTGTGTCAGTTAG |
| loxP-G418-loxP NheI antisense | TGCTTAGCTAGCTGATAACTTCGTATAATGTATGCTATACGAAGTTATTTTATTCTGT |
| G418 cassette for C-terminal KI: | |
| G418 NotI sense | cgtcgGCGGCCGCcctgaggcggaaagaaccagctgtggaatgtgtgtcagttag |
| G418 EcoRI antisense | tcatgGAATTCtttattctgtctttttattgccgtc |
| Puro cassette for C-terminal KI: | |
| Puro NotI sense | GCTCAGCGGCCGCCTGTGGAATGTGTGTCAGTTAGGGT |
| Puro EcoRI antisense | CAACTGAATTCCAGACATGATAAGATACATTGATGA |
| SNAP-tag for C-terminal KI: | |
| SNAP BamHI sense | ctgatGGATCCGACAAAGACTGCGAAATGAAGCGCA |
| SNAP V5 NheI antisense | TACAAGCTAGCTCAGGTGCTGTCCAGGCCCAGCAGGGGGTTGGGGATGGGCTTGCCACCCAGCCCAGGCTTGCCCAGTCTG |
| HaloTag for C-terminal KI: | |
| Halo BamHI sense | ctgatGGATCCATGGCAGAAATCGGTACTGGCTTTC |
| Halo NheI antisense | TGCTTAGCTAGCGCCGGAAATCTCGAGCGTCGACAGC |
| Halo ALFA NheI antisense | TGCCTGCTAGCTCATGGCTCGGTCAGTCTTCTTCTCAGCTCCTCCTCCAGTCTGCTTGGGCCGGAAATCTCGAGCGTGGACAGC |
| eGFP-tag for ARF1 KI: | |
| eGFP NheI sense | TATCTGCTAGCATGGTGAGCAAGGGCGAGGAGCTGTTCA |
| eGFP BamHI antisense | TATCTGGATCCTCACTTGTACAGCTCGTCCATGCCGAGA |
| PolyA fragment for C-terminal KI: | |
| PolyA NheI sense | AGTTCGCTAGCCCGCGACTCTAGATCATAATCAGC |
| PolyA NotI antisense | tgcacGCGGCCGCcttacaatttacgccttaagataca |
| ARF1-SNAP-V5-PolyA-Puro^R^ HR plasmid: | |
| SNAP NheI sense | TTCAGTGCTAGCATGGACAAAGACTGCGAAATGAAGCGCA |
| SNAP-V5 antisense | TACAAGCTAGCTCAGGTGCTGTCCAGGCCCAGCAGGGGGTTGGGGATGGGCTTGCCACCCAGCCCAGGCTTGCCCAGTCTG |
| Puro BamHI antisense | AATTTGGATCCCAGACATGATAAGATACATTGATGAGTTTG |
| Hygromycin^R^ HR plasmid: | |
| Hygromycin ClaI sense | AATTATCGATATGaaaaagcctgaactcaccgcgacgt |
| Hygromycin EcoRI antisense | AATTGAATTCtaagatacattgatgagtttggaca |

**Supplementary Table 2: List of guide RNA sequences and used vectors**

| Targeted locus | Guide RNA sequence  (PAM side underlined) | Vector |
| --- | --- | --- |
| KI guides: | | |
| AP1M1 (+ strand) (Gene ID 8907) | 5´-CAGCCAACACCCCGGCCTCGGGG-3’ | pX459 |
| AP2M1 (+ strand) (Gene ID 1173) | 5´- ACTCGCTGCTAGCTGCCACTAGG-3’ | pX459 |
| AP3M1 (- strand) (Gene ID 26985) | 5´- TGGAAAACAAACTGGTCCTGAGG-3’ | pX459 |
| AP4M1 (+ strand) (Gene ID 9179) | 5´- GATCTGAGGCTCCCCAAACGAGG-3’ | pX330 |
| AP1G1 (+ strand) (Gene ID 164) | 5´- TCTTTATCCCACTCAATCAAAGG-3’ | pX330 |
| AP3D1 (- strand) (Gene ID 8943) | 5´- CCCTGGGTACGTGCTCCGCGGGG-3’ | pX330 |
| CLTA (+ strand) (Gene ID 1211) | 5’- ATGGCTGAGCTGGATCCGTTCGG-3’ | pX330 |
| ARF1 (+strand) (Gene ID 375) | 5’- CCACATGAGAGTAAAGCAGAGGG-3’ | pX330 |
| Rab11A (-strand) (Gene ID 8766) | 5’- CGTCGCGGGTGCCCATTGCGCGG-3’ | pX459 |
| Rab6A (+strand) (Gene ID 5870) | 5’- AGTTCCACAATGTCCACGGGCGG-3’ | pX459 |
| Rab7A (+strand) (Gene ID 7879) | 5’- TAGTTTGAAGGATGACCTCTAGG-3’ | pX330 |
| SNX1 (+strand) (Gene ID 6642) | 5’- TGGAAGAAGATGGCGTCGGGTGG-3’ | pX330 |
| KO guides: | | |
| AP1M1 (+ strand, Exon 2) (Gene ID 8907) | 5’- CCGGAACTACCGTGGCGACGTGG-3’ | pX459 |
| AP1M1 (+ strand, Exon 5) (Gene ID 8907) | 5’- ACCAGCCACCGTCACCAACGCGG-3’ | pX459 |

**Supplementary Table 3: CRISPR cell lines and selection**

| Cell line | Base cell line | Plasmids used | Selection |
| --- | --- | --- | --- |
| HAP1 ARF1^EN^-Halo-PolyA-G418^R^/ SNAP-CLCa^EN^ | HAP1 ARF1^EN^-Halo-PolyA-G418^R 6^ | loxP-Puro^R^-loxP-SNAP-CLCa HR plasmid, CLCa guide | Puromycin |
| HAP1 AP1µA^EN^-Halo-PolyA-G418^R^ | HAP1 WT | AP1µA-Halo-PolyA-G418^R^ HR plasmid, AP1M1 guide | G418 |
| HeLa ARF1^EN^-Halo-PolyA-G418^R^/ AP1µA^EN^-SNAP-V5-PolyA-Puro^R^ | HeLa ARF1^EN^-Halo-PolyA-G418^R 7^ | AP1µA-SNAP-V5-PolyA-Puro^R^ HR plasmid, AP1M1 guide | Puromycin |
| HeLa ARF1^EN^-Halo-PolyA-G418^R^/ AP3µA^EN^-SNAP-V5-PolyA-Puro^R^ | HeLa ARF1^EN^-Halo-PolyA-G418^R^ | AP3µA-SNAP-V5-PolyA-Puro^R^ HR plasmid, AP3M1 guide | Puromycin |
| HeLa ARF1^EN^-eGFP | HeLa WT | ARF1-eGFP HR plasmid, ARF1 guide | FACS sorting (2% brightest) |
| HeLa ARF1^EN^-eGFP/AP3µA^EN^-SNAP-V5-PolyA-Puro^R^ | HeLa ARF1^EN^-eGFP | AP3µA-SNAP-V5-PolyA-Puro^R^ HR plasmid, AP3M1 guide | Puromycin |
| HeLa ARF1^EN^-eGFP/AP3µA^EN^-SNAP-V5-PolyA-Puro^R^/AP1µA^EN^-Halo-ALFA-PolyA-G418^R^ | HeLa ARF1^EN^-eGFP/AP3µA^EN^-SNAP-V5-PolyA-Puro^R^ | AP1µA-Halo-ALFA-PolyA-G418^R^ HR plasmid, AP1M1 guide | G418 |
| HeLa ARF1^EN^-eGFP/AP3µA^EN^-SNAP-V5-PolyA-Puro^R^ /Halo-CLCa^EN^ | HeLa ARF1^EN^-eGFP/AP3µA^EN^-SNAP-V5-PolyA-Puro^R^ | loxP-G418^R^-loxP-Halo-CLCa HR plasmid, CLCa guide | G418 |
| HeLa ARF1^EN^-eGFP/AP1µA^EN^-SNAP-V5-PolyA-Puro^R^ | HeLa ARF1^EN^-eGFP | AP1µA-SNAP-V5-PolyA-Puro^R^ HR plasmid, AP1M1 guide | Puromycin |
| HeLa ARF1^EN^-eGFP/AP1µA^EN^-SNAP-V5-PolyA-Puro^R^/Halo-CLCa^EN^ | HeLa ARF1^EN^-eGFP/AP1µA^EN^-SNAP-V5-PolyA-Puro^R^ | loxP-G418^R^-loxP-Halo-CLCa HR plasmid, CLCa guide | G418 |
| HeLa ARF1^EN^-Halo-PolyA-G418^R^/ SNAP-CLCa^EN^ | HeLa ARF1^EN^-Halo-PolyA-G418^R^ | loxP-Puro^R^-loxP-SNAP-CLCa HR plasmid, CLCa guide | Puromycin |
| HeLa ARF1^EN^-Halo-PolyA-Puro^R^/ AP1γA^EN^-SNAP-V5 | HeLa ARF1^EN^-Halo-PolyA-G418^R^ | AP1γA-SNAP-V5-PolyA-Puro^R^ HR plasmid, AP1G1 guide | Puromycin |
| HeLa ARF1^EN^-Halo-PolyA-G418^R^/ AP3δA^EN^-SNAP-V5-PolyA-Puro^R^ | HeLa ARF1^EN^-Halo-PolyA-G418^R^ | AP3δA-SNAP-V5-PolyA-Puro^R^ HR plasmid, AP3D1 guide | Puromycin |
| HeLa AP1µA^EN^-Halo-PolyA-G418^R^ | HeLa WT | AP1µA-Halo-PolyA-G418^R^ HR plasmid, AP1M1 guide | G418 |
| HeLa AP3µA^EN^-Halo-ALFA-PolyA-G418^R^ | HeLa WT | AP3µA-Halo-ALFA-PolyA-G418^R^ HR plasmid, AP3M1 guide | G418 |
| HeLa APµ1A^EN^-Halo-PolyA-G418^R^/ AP1γA^EN^-SNAP-V5 | HeLa AP1µA^EN^-Halo-PolyA-G418^R^ | AP1γA-SNAP-V5-PolyA-Puro^R^ HR plasmid, AP1G1 guide | Puromycin |
| HeLa AP3µA^EN^-Halo-ALFA-PolyA-G418^R^/ AP3δA^EN^-SNAP-V5 | HeLa AP3µA^EN^-Halo-ALFA-PolyA-G418^R^ | AP3δA-SNAP-V5-PolyA-Puro^R^ HR plasmid, AP3D1 guide | Puromycin |
| HeLa Halo-Rab7A^EN^ | HeLa WT | loxP-G418^R^-loxP-Halo-Rab7A HR, Rab7A guide | G418 |
| HeLa ARF1^EN^-SNAP-V5-PolyA-Puro^R^ /Halo-Rab7A^EN^ | HeLa Halo-Rab7A^EN^ | ARF1-SNAP-V5-PolyA-Puro^R^ HR plasmid, ARF1 guide | Puromycin |
| HeLa Halo-Rab11A^EN^ | HeLa WT | loxP-G418^R^-loxP-Halo-Rab11A HR, Rab11A guide | G418 |
| HeLa ARF1^EN^-SNAP-V5-PolyA-Puro^R^ /Halo-Rab11A^EN^ | HeLa Halo-Rab11A^EN^ | ARF1-SNAP-V5-PolyA-Puro^R^ HR plasmid, ARF1 guide | Puromycin, FACS sorting (5% brightest) |
| HeLa Halo-Rab11A^EN^ / ARF1^EN^-mStayGold-PolyA-Hygromycin^R^ | HeLa Halo-Rab11A^EN^ | ARF1-mStayGold-PolyA-Hydromycin HR, ARF1 guide | Hygromycin |
| HeLa 3xALFA-Halo-Rab6A^EN^ | HeLa WT | loxP-G418^R^-loxP-3xALFA-Halo-Rab6A HR, Rab6A guide | G418 |
| HeLa ARF1^EN^-SNAP-V5-PolyA-Puro^R^ /3xALFA-Halo-Rab6A^EN^ | HeLa Halo-3xALFA-Rab6A^EN^ | ARF1-SNAP-V5-PolyA-Puro^R^ HR plasmid, ARF1 guide | Puromycin |
| HeLa ARF1^EN^-SNAP-V5-PolyA-Puro^R^ /3xALFA-Halo-Rab6A^EN^ / AP1µA^EN^-mStayGold-PolyA-Hygromycin^R^ | HeLa ARF1^EN^-SNAP-V5-PolyA-Puro^R^ /3xALFA-Halo-Rab6A^EN^ | AP1µA-mStayGold-PolyA-Hydromycin HR, AP1M1 guide | Hygromycin |
| HeLa 3xALFA-Halo-SNX1^EN^ | HeLa WT | loxP-G418^R^-loxP-3xALFA-Halo-SNX1 HR, SNX1 guide | G418 |
| HeLa ARF1^EN^-SNAP/3xALFA-Halo-SNX1^EN^ | HeLa 3xALFA-Halo-SNX1^EN^ | ARF1-SNAP-loxP-Puro^R^-loxP HR plasmid, ARF1 guide | Puromycin |
| HeLa SNAP-CLCa^EN^/Halo-Rab11A^EN^ | HeLa Halo-Rab11A^EN^ | loxP-Puro^R^-loxP-SNAP-CLCa HR, CLCa guide | Puromycin |
| HeLa AP1µA KO ARF1^EN^-Halo-PolyA-G418^R^/SNAP-CLCa^EN^ | HeLa ARF1^EN^-Halo-PolyA-G418^R^/SNAP-CLCa^EN^ | AP1M1 KO guide 1&2 | Puromycin + single cell clone |
| HeLa AP1µA KO ARF1^EN^-SNAP-PolyA-Puro^R^/Halo-Rab11A^EN^ | HeLa ARF1^EN^-SNAP-PolyA-Puro^R^/Halo-Rab11A^EN^ | AP1M1 KO guide 1&2 | Puromycin + single cell clone |
| Jurkat ARF1 ^EN^-SNAP-PolyA-G418^R^/Halo-CLCa^EN^ | Jurkat WT | ARF1-SNAP-PolyA-G418^R^ HR plasmid, ARF1 guide, loxP-G418^R^-loxP-Halo-CLCa HR plasmid, CLCa guide | G418 |

**Supplementary Table 4: Antibodies and conjugates**

| Antibody/Conjugate | Supplier | Used concentration |
| --- | --- | --- |
| β-Actin Mouse (8H10D10) | Cell Signaling | 1:1000 for WB |
| Anti-AP1M1 Rabbit (ab230273) | Abcam | 1:1000 for WB |
| Anti-CHC Mouse (NB300-613) | Novus Biologicals | 1:1000 for IF |
| Anti-HaloTag Mouse (G921) | Promega | 1:2000 for WB |
| Anti-SNAP Rabbit (A00684) | Genscript | 1:1000 for WB |
| Transferrin-AF488 (T13342) | Thermo Fisher Scientific | 25 µg/ml |
| Holo-Transferrin (T0665) | Sigma Aldrich | 0,5 mg/ml |

**Supplementary Table 5: CellProfiler script used to quantify the overlap between ARF1 compartments and different post-Golgi markers**

| **ARF1+Rab5/7/11 or SNX1** | | |
| --- | --- | --- |
|  | **ARF1** | **Marker** |
| Size restriction | Pixels: 4-50 | Pixels: 4-30 |
|  | Discard objects outside the range and touching the border of the image | |
| Threshold | Adaptive, Robust background (0.05 lower and upper outlier format), averaging method: mean, variance method: standard deviation, number of deviations = 0, threshold smoothing scale = 1, threshold correction factor = 1.2, lower and upper bounds on threshold: 0.0 and 1.0, size adaptation window: 2  Method to distinguish clumped objects: none | Adaptive, Robust background (0.05 lower and upper outlier format), averaging method: mean, variance method: standard deviation, number of deviations = 0, threshold smoothing scale = 1, threshold correction factor = 1.3, lower and upper bounds on threshold: 0.0 and 1.0, size adaptation window: 2  Method to distinguish clumped objects: Intensity |
|  | Fill holes in identified objects: After both thresholding and declumping | |
| **ARF1+Rab6 or ARF1 (positive control)** | | |
|  | **ARF1** | **Marker** |
| Size restriction | Pixels: 4-50  Discard objects outside the range and touching the border of the image | |
| Threshold | Adaptive, Robust background  0.05 lower and upper outlier format, averaging method: mean, variance method: standard deviation, number of deviations = 0, threshold smoothing scale = 1, threshold correction factor = 1.2, lower and upper bounds on threshold: 0.0 and 1.0, size adaptation window: 2 | Adaptive, Robust background  0.05 lower and upper outlier format, averaging method: mean, variance method: standard deviation, number of deviations = 0, threshold smoothing scale = 1, threshold correction factor = 1.15, lower and upper bounds on threshold: 0.0 and 1.0, size adaptation window: 2 |
|  | Method to distinguish clumped objects: none  Fill holes in identified objects: After both thresholding and declumping | |

**Supplementary Table 6: CellProfiler script used to determine exact sorting step ARF1 compartments facilitate endocytic recycling of Tfn in relation to other endosomal markers**

| **Tfn + ARF1/Rab6** | | |
| --- | --- | --- |
|  | **Marker** | **Tfn** |
| Smooth | Artificial diameter Gaussian filter: 3 | |
| Size restriction | Pixels: 2-35  Discard objects outside the range and touching the border of the image | |
| Threshold | Adaptive Otsu  Two classes, threshold smoothing scale = 1, threshold correction factor = 0.7, lower and upper bounds on threshold: 0.0 and 1.0, size adaptation window: 4  Method to distinguish clumped objects: none | Adaptive Otsu  Two classes, threshold smoothing scale  = 1, threshold correction factor =  1.1, lower and upper bounds on threshold: 0.0 and 1.0, size adaptation window: 4  Method to distinguish clumped objects: none |
|  | Fill holes in identified objects: After both thresholding and declumping | |
| **Tfn+Rab5/11** | | |
|  | **Marker** | **Tfn** |
| Smooth | Artificial diameter Gaussian filter: 3 | |
| Size restriction | Pixels: 2-35  Discard objects outside the range and touching the border of the image | |
| Threshold | Adaptive Otsu  Two classes, threshold smoothing scale = 1, threshold correction factor = 1.1, lower and upper bounds on threshold: 0.0 and 1.0, size adaptation window: 4  Method to distinguish clumped objects: none | |
|  | Fill holes in identified objects: After both thresholding and declumping | |
